# Machine learning-guided design of human gut microbiome dynamics in response to dietary fibers

**DOI:** 10.1101/2025.03.03.641341

**Authors:** Bryce M. Connors, Jaron Thompson, Ophelia S. Venturelli

## Abstract

Dietary fibers are key modulators of human gut microbiome dynamics and functions, yet we lack the ability to predict community dynamics and functions in response to fibers. We integrate machine learning, Bayesian optimization, and high-throughput community construction to investigate how dietary fibers shape health-relevant functions of human gut microbial communities. To efficiently navigate the landscape of fiber-microbiome interactions, we implemented a design-test-learn cycle to identify fiber-species combinations that maximize a multi-objective function capturing beneficial community properties. Our model-guided approach revealed a highly butyrogenic and robust ecological motif characterized by the copresence of inulin, *Bacteroides uniformis*, and *Anaerostipes caccae* and a higher-order interaction with *Prevotella copri*. In germ-free mice, model-designed species-fiber combinations impacted colonization, community diversity and short-chain fatty acid profiles. In sum, our work establishes a novel framework for designing microbial communities with desired functions in response to key nutrients such as dietary fibers.

## INTRODUCTION

The gut microbiome is a complex and dynamic ecosystem that plays a fundamental role in host health, influencing metabolism, immune function, and disease susceptibility^1–4^. As a primary energy source for gut bacteria, dietary fibers are a major driver of the dynamics of the human gut microbiome and substantially impact health-relevant functions of this complex system^5,6^. Gut bacteria harbor an arsenal of specialized enzymatic machinery broadly classified as carbohydrate active enzymes (CAZymes) organized into polysaccharide utilization loci (PULs) for utilizing dietary fibers with diverse chemical and physical properties^7^. The presence of dietary fibers yields substantial shifts in the composition and function of the microbiome that are largely unpredictable^8,9^. Developing the capability to uncover the web of interactions linking dietary fibers and human gut bacteria could guide novel precision interventions that steer microbiomes to desired metabolic states for preventing or treating multiple human diseases.

The production of certain short-chain fatty acids, notably butyrate, by human gut bacteria is strongly associated with a healthy gut state^10,11^. Butyrate serves as the primary energy source for colonocytes, activates anti-inflammatory signaling pathways, and bolsters gut barrier integrity^12–15^. Consistent with these beneficial functions, depletion of butyrate producers has been associated with multiple human diseases^16,17^. Initial breakdown of polymeric fiber structures is performed by primary fermenters possessing specific PULs, harbored frequently in Bifidobacteria and Bacteroides^18,19^. Shorter chain oligosaccharides released during the breakdown process, as well as organic acid fermentation byproducts, can fuel the growth and metabolism of butyrate producing organisms, predominantly in Clostridia^20^. Taxonomically diverse microbiomes contain a wide range of metabolic pathways for degrading fibers. In addition, diverse microbiomes display enhanced colonization resistance to invading species (e.g. pathogens) by blocking access to ecological niches^21,22^. While dietary fibers and the composition of the gut microbiome play a critical role in shaping host health, the interactions between fibers and beneficial microbes such as butyrate producing bacteria remain poorly understood. The bottom-up construction of synthetic microbial communities is a powerful approach for gaining deeper insights into the interactions between dietary fibers and bacteria^23–26^. While interactions between primary degraders and butyrate producers have been studied in low richness synthetic communities in response to individual fibers, the types of interactions shaping butyrate production and diversity in larger communities in the presence of combinations of fibers has not yet been resolved^27,28^.

Considering only the presence or absence of species and fibers, the number of conditions grows exponentially, rendering an exhaustive characterization intractable using high-throughput methods. An effective strategy to rationally navigate vast experimental design spaces is to leverage data-driven computational models that can learn to predict microbial community functions from experimental data that sparsely sample the design space^23,24^. While the complexity of microbial consortia limits the use of mechanistic models based on first principles, machine learning (ML) models have proven to be effective tools to predict microbial community functions from training data^29^. Machine learning models that are designed to model sequential data such as recurrent neural networks (RNNs) are particularly well suited to predict the dynamics of microbial species and metabolites^30,31^. While standard RNNs can make physically unrealistic predictions such as the emergence of species that were not inoculated and negative species abundances or metabolite concentrations, the physically constrained Microbiome RNN (MiRNN) can overcome these limitations^32^. Once such models are trained and validated on experimental data, interpretable machine learning methods such as local interpretable model-agnostic explanations (LIME) and Shapley additive explanations (SHAP) can be used to derive biological insights by explaining model predictions^33,34^.

Bayesian optimization is an experimental design algorithm that leverages model prediction uncertainty to select conditions expected to simultaneously improve learning (exploration) and achieve specific goals (exploitation). By focusing on exploration of new conditions where predictions are uncertain as well as exploitation of conditions likely to yield the best results, Bayesian optimization efficiently fills knowledge gaps and optimizes system outputs^35^. Bayesian optimization has proven to be an effective experimental design strategy in protein engineering, chemical synthesis, and materials discovery, but has not yet been experimentally validated for the design of defined microbial consortia^36,37^.

To elucidate fundamental, quantitative principles by which fibers shape community composition and functions, we considered an experimental system of 15 diverse and prevalent human gut species and six dietary fibers that vary in chemical and physical properties^38^. Leveraging a tailored Bayesian recurrent neural network (MiRNN), we demonstrated the ability to accurately predict species growth and metabolite production as a function of inoculated species and fiber combinations. Using Bayesian optimization based experimental design and high-throughput community assembly, we identified species-fiber combinations that maximized health-relevant community properties including ecological diversity, persistence of composition over time (i.e. minimal variation in composition), and butyrate production. This task was achieved over three DTL cycles wherein the MiRNN was updated at each stage with measurements of species abundances and metabolites concentrations. Explainable machine learning was used to identify underlying ecological principles arising from interactions among combinations of fibers and species that govern the variation in butyrate production. We identified a third-order interaction where the combination of a fiber, inulin, a primary fiber degrader, *Bacteroides uniformis*, and a butyrate producer, *Anaerostipes caccae*, produced greater levels of butyrate than any combination of its components. We also found a fourth-order interaction where *Prevotella copri* or *B. uniformis* individually promoted butyrate production in microbial communities. However, when both species are present together, butyrate production was significantly reduced. Finally, we investigated colonization, short-chain fatty acid profiles and community dynamics of model-designed combinations of fibers and species in the mammalian gut using germ-free mice. Our integrated experimental and computational framework for exploring the functional landscapes of human gut communities shaped by combinations of dietary fibers provides a foundation for the rational design of precision microbiome interventions to steer these systems to desired states.

## RESULTS

### Overview of framework for the design of community dynamics and functions

A key knowledge gap is whether the compositional and functional landscapes of complex microbial communities can be quantitatively predicted in response to combinations of fibers. We explored the mappings between combinations of fibers and species and community assembly and functions by integrating high-throughput experiments and computational modeling. For the target functions of this system, we focused on community properties that are linked with the healthy gut state: butyrate production, Shannon diversity, and persistence in composition over time. Butyrate production is a signature of a healthy microbiome state whereas a reduction in butyrate has been linked to dysbiosis in inflammatory bowel disease and antibiotic induced microbiome disruption^39–41^. A survey of longitudinal studies of the human gut microbiome revealed consistent qualitative stability over timescales of months or years in healthy adults^42^. Stability of the microbiome is generally quantified by a measure of compositional similarity between time points^42,43^. Instability and reduced diversity has been associated with a variety of negative health outcomes including inflammatory bowel disease, metabolic disease, and COVID-19^44–47^. The ability to understand and design these functions constitutes an important step towards realizing the potential for rational design of health-beneficial microbiome interventions.

Our microbial design space consisted of 15 prevalent human gut isolates spanning the major human gut phyla including Firmicutes, Bacteroides, and Actinobacteria (**Fig. 1a**). The community contains the health-beneficial butyrate producing strains *Anaerostipes caccae, Coprococcus comes, Eubacterium rectale, Faecalibacterium prausnitzii, Roseburia intestinalis*^48,4950^. Our resource design space consisted of six health-relevant fibers with different chemical structures and selectivities^47^. Inulin and acacia gum (arabinogalactan is a major component) have been demonstrated to increase both butyrate production in human subjects and health-relevant Bifidobacterium *in vitro*^51–53^. Pectin is present in fruit and has been a component of the human diet for over hundreds of thousands of years^54^. Pectin has been shown to modulate microbial butyrate production via cross feeding and protect intestinal epithelial permeability as well as increase cecal short-chain fatty acid concentrations in murine disease models^55,56^. Xylan, an abundant component of cereal grains, has been shown to exhibit health-promoting properties^57^. Starch constitutes around 25% of caloric intake, and while a substantial fraction is digested by the host enzymes including amylase, resistant starch reaches the colon and is an influential factor for microbial short-chain fatty acid production^58,59^.

**Figure 1.**
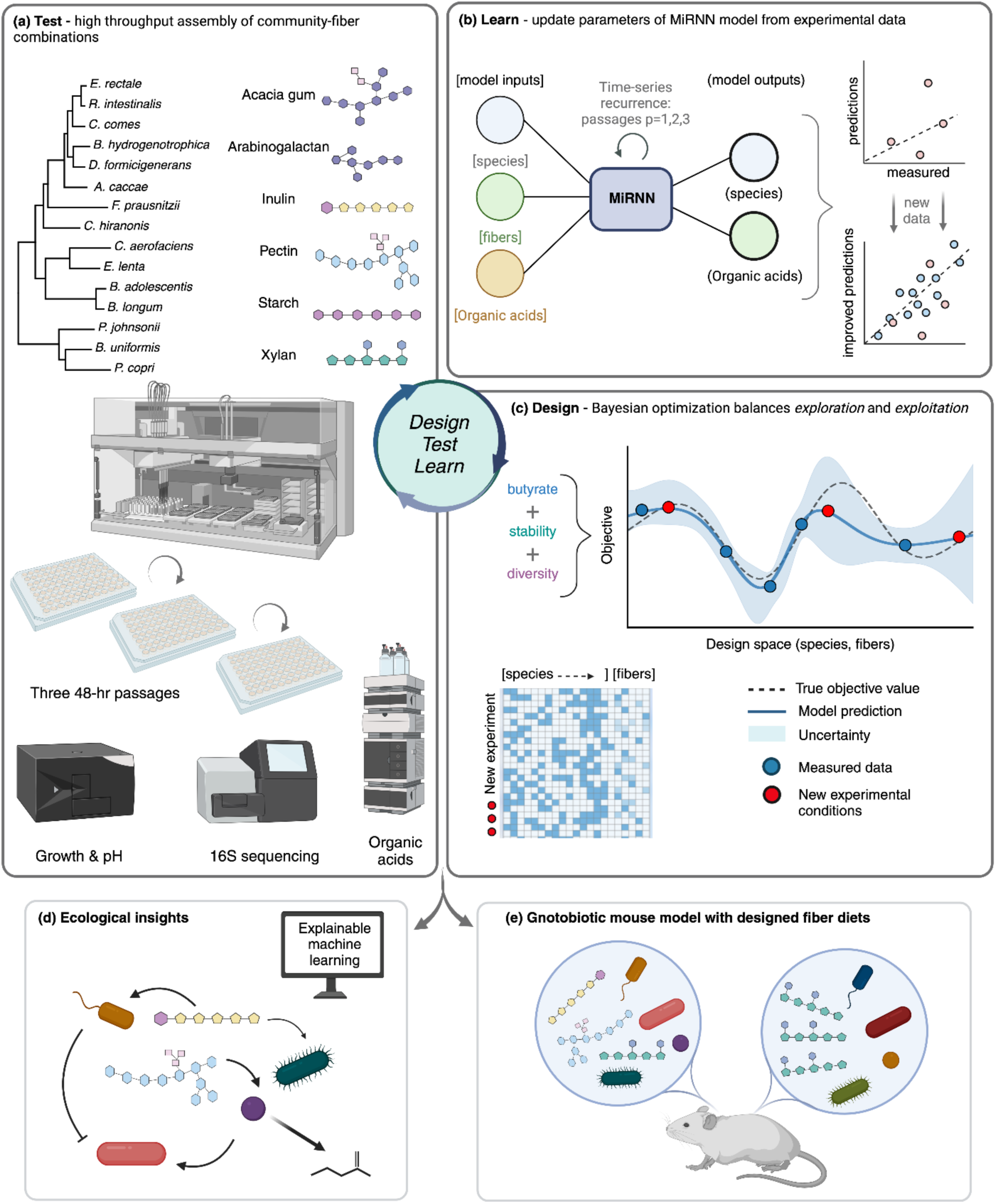
Exploring a large species-fiber experimental design landscape with Bayesian optimization. (**a)** The design space is comprised of fifteen phylogenetically and functionally diverse commensal species and six fibers totaling over two million possible combinations (2^15^ * 2^6^). Species and fibers are arrayed in high-throughput using a liquid handling robot and cultured anaerobically for three 48-hour passages. Growth, community composition, and organic acids are assayed via OD600, 16S sequencing, and HPLC, respectively. Each of the three DTL cycle experiments evaluates 288 species-fiber combinations over a time-series of three passages via 50-fold dilution into fresh media and culturing until approximately stationary phase. (**b)** A tailored Bayesian recurrent neural network model (MiRNN) predicts timeseries species and metabolite abundances from previous-passage experimental data and fiber initial conditions. (**c)** A Bayesian optimization experimental design algorithm leverages model predictions and uncertainty to design new batches of experimental conditions based on a criterion balancing both model predictive improvement (exploration) and objective improvement (exploitation). The design objective is a weighted sum of three human health-relevant community attributes: Shannon diversity, butyrate production, and temporal stability (Methods). The new batch of designed species-fiber conditions is evaluated experimentally in a subsequent iteration of the DTL cycle. (**d)** Explainable machine learning methods were applied to the trained MiRNN to derive ecological insights relating species and fibers to community functions. (**e)** Optimized communities were characterized in a gnotobiotic mouse model to investigate colonization ability, community diversity and metabolite profiles.

The relationship between system variables and total combinations is exponential in nature^60^. Our 21 fiber and species variables yield a design space of over 2,000,000 species-fiber combinations, a magnitude for which exhaustive characterization is experimentally intractable using high-throughput experimental techniques. However, we hypothesized that the underlying rules leading to these functional and compositional landscapes could be learned from a much smaller subset of informative conditions, obviating the need for exhaustive characterization. Learning these underlying rules is an important step towards understanding how distinct systems-level community properties like butyrate production and ecological diversity emerge from interactions between specific combinations of fiber and bacterial species. We therefore used a Bayesian optimization approach (**Fig. 1b**) to design experiments composed of an experimentally tractable subset of species and fiber combinations that simultaneously maximize a design objective (exploitation) and information content (exploration). For studying fiber-species combinations, the MiRNN can simulate the outcome of previously untested experimental conditions to identify those which balance model prediction uncertainty with desired community dynamics and functions^61^. Rationally designed experiments can then be tested using high-throughput liquid handling robotics, generating highly informative data to further refine the model. This iterative process of designing new species-fiber combinations, testing these designs, and updating the model with experimental data embodies an active “design-test-learn” (DTL) approach to microbial community optimization^62^.

Functions that incorporate both exploitation and exploration to quantify the utility of candidate experimental designs are called acquisition functions^63^. Our design objective is a weighted sum of Shannon diversity, butyrate production, and temporal persistence of composition. The information content of an experimental design is quantified by the expected information gain (EIG), which measures the degree to which the expected data constrains the parameter distribution of the model (**Methods**)^64^. The parameter distribution conditioned on experimental data (i.e. the posterior) is approximated as a Gaussian distribution. The mean of the Gaussian reflects the magnitude or importance of the parameter, and the variance reflects the probabilistic uncertainty associated with inference. The parameter distribution was inferred from time-series experimental data using an expectation-maximization (EM) algorithm (**Methods**)^65^.

To predict microbial communities, we used the MiRNN (**Fig 1c**)^66^. In general, RNNs are flexible machine learning models that can be adapted to predict a variety of input and output variables over discrete time intervals. Passaging experiments, in which communities are iteratively cultured for a period of time and then used to inoculate a new culture, have been used to study microbial communities over longer timescales^67,68^. The MiRNN is thus highly fit-for-purpose because its discrete time interval architecture mirrors the discrete time-series measurements of passaging experiments. By contrast, continuous time models such as ordinary differential equations require simulation of the dilution step at each passage and are more computationally expensive due to the need for numerical integration. The tailored structure of the MiRNN enables the prediction of bacterial species and metabolite abundances over time given an initial condition. The MiRNN also incorporates constraints to prevent the model from predicting negative species abundances or the emergence of species that were not inoculated.

Bayesian modeling and optimization integrates naturally into the DTL cycle paradigm^61^. Although historically associated with the fields of protein and metabolic engineering, Bayesian optimization has not yet been applied to design synthetic microbial communities *in vitro*^69,70^. The success of this DTL approach is rooted in the ability to iteratively navigate immense design spaces and functional landscapes (e.g. sequence space). The “design” phase consisted of leveraging the MiRNN informed by experimental observations via Bayesian optimization to select a set of informative, high-objective community-fiber conditions for experimental characterization. In the “test” phase, these designed community-fiber combinations were assembled in high-throughput using a liquid-handling robot housed inside an anaerobic chamber and cultured for three 48-hour passages (approximately stationary phase, **Supplementary Figure 1a**). Growth, species composition, and organic acid production were measured via OD600, 16S sequencing, and HPLC at the end of the passages (**Methods**). In the “learn” phase, the new data was incorporated to train the MiRNN.

### Bayesian optimization over the species-resource-function landscape

To provide an initial data set for model training that accounted for the presence of all species and fibers, we characterized the full community and communities of all species except one (leave-one-out) communities cultured in the presence of individual fibers or single-fiber dropout media. Single species deletion communities can provide insights into the effects of individual species on community functions^71,72^. Additionally, we performed time-series measurements of the growth of single species in the presence of individual fibers, single-fiber dropouts, and no-fiber media. In total, this initial data set comprised 387 community-fiber combinations (DTL 0). We used a defined media that supports the growth of diverse human gut bacteria supplemented with fibers (5 grams per liter)^50^. *Anaerostipes caccae, Bifidobacterium adolescentis, Bifidobacterium longum, Bacteroides uniformis, Coprococcus comes*, and *Prevotella copri* displayed substantial growth (OD600 > 0.5) in the presence of at least one fiber, while only *B. uniformis* and *P. copri* displayed substantial growth in multiple fibers (**Supplementary Figure 1a**). In the absence of fiber, growth was not observed for any species.

Overall, monoculture growth was relatively sparse with only 37 of 90 single species, single fiber conditions yielding a substantial increase (50-fold) in OD600 compared to the inoculation density (**Supplementary Figure 1b**). By contrast, the communities yielded more than a 50-fold increase in OD600 in 89 of 90 leave-one-out conditions in single fibers. The significant increase in growth for communities compared to monoculture conditions across fibers suggests that positive inter-species interactions enhanced growth (Mann-Whitney U-test p-values < 1e-7) (**Supplementary Figure 1b**). The single fiber communities were diverse, with their mean and maximum Shannon diversities of 1.73 and 2.26 corresponding to 64% and 83% of the maximum Shannon diversity for a 15-member community. The highest diversity community was observed in a single-fiber medium (pectin). The mean Shannon diversity of communities in the presence of five fibers (single fiber dropouts) was approximately 10% higher than the diversity of communities cultured in the presence of individual fibers (p-value < 1e-10, Mann-Whitney U test, **Supplementary Figure 1d**). These data demonstrate that a single fiber type as the sole carbon source in a defined medium can support high ecological diversity.

The initial dataset of 387 community-fiber combinations (DTL0) was used to train the MiRNN model (MiR0) (**Supplementary Table 1**). As the dataset consisted of either high or low richness communities representing the extremes of the species design space, the model will have limited prediction capabilities. We therefore implemented a Bayesian experimental design algorithm (pure exploration) selecting 288 new species-fiber combinations for the design of the first model-guided experiment (DTL1). After training the model on DTL1 data, we designed communities to maximize a weighted combination of butyrate production, Shannon diversity, and compositional persistence across passages (a proxy for stability) (**Fig. 2a**, **Methods**). To improve model prediction performance and seek conditions that optimized our design objective, we performed two additional DTL cycles (DTL2 and 3) using Bayesian optimization algorithms (**Methods**) to select 288 new species-fiber combinations that favored both improvement in model prediction performance (exploration) and maximization of the design objective (exploitation). Finally, we performed a validation experiment with 4 replicates of the top 8 conditions from DTL 3 (DTL3 validation) and a final DTL cycle with 4 replicates of 8 conditions selected using pure exploitation of the design objective (DTL4). The median objective value increased with each DTL cycle (0.32, 0.44, 0.91, and 1.10), and a substantial improvement in the maximum objective value was observed from DTL2 to DTL3 (1.10 to 1.36, 24% improvement). The maximum objective value in DTL4 was only slightly higher than the maximum value observed in DTL3 at 1.38, suggesting that further DTL cycles would not lead to substantial improvements in the objective (**Fig. 2b**).

**Figure 2.**
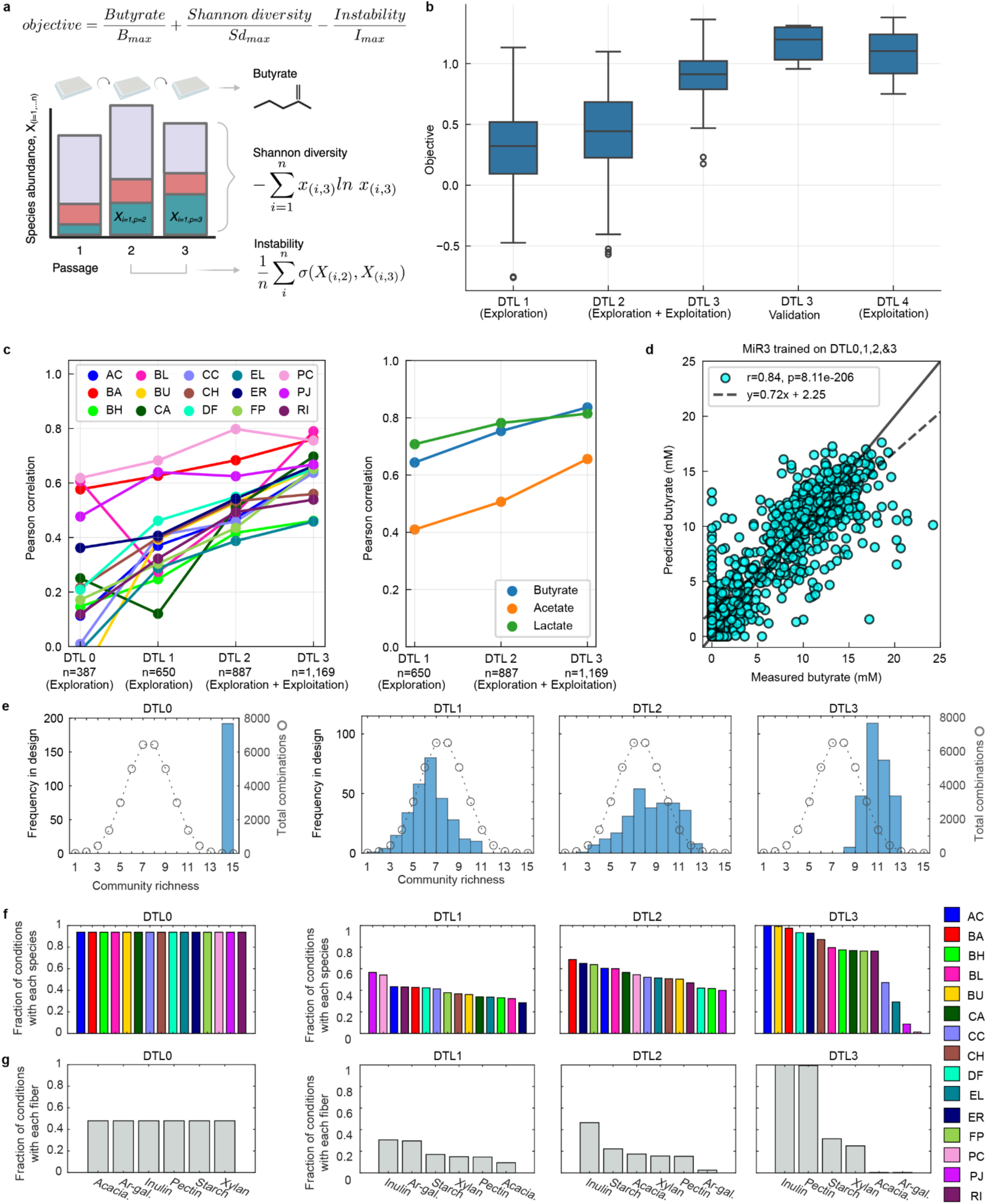
Model predictive performance across Design-Test-Learn cycles. **(a)** Overview of design objective function quantifying butyrate, diversity, and temporal stability of species-fiber systems. Butyrate and Shannon diversity are quantified at the endpoint of the 3^rd^ passage. As a proxy for stability, instability is quantified as the average standard deviation of each species abundance between passage two and three. These quantities are normalized by their respective maximum values and summed to a scalar-valued objective function. (**b)** Boxplot of objective values for experiments DTL1 (n=288), DTL2 (n=288), DTL3 (n=288), DTL3 validation (n=8), and DTL4 (n=8). The objective is computed as the normalized sum of Shannon diversity, butyrate, and stability after the 3^rd^ passage (Methods). Center line, box edges, and whiskers indicate median, quartiles, and quartile plus 1.5 times interquartile range, respectively. Diamonds indicate values outside of the range spanned by the whiskers. (**c)** Line plots of model predictive performance of the MiRNN for the 15 species and 3 metabolites. Predictive performance is quantified as the Pearson correlation coefficient between predicted and measured values. A cross-validation based approach evaluates models trained on a particular subset of predicted and DTL1) for prediction performance on the entire dataset (Methods). (**d)** Scatter plot of model predictions vs. measured butyrate concentration after three DTL cycles. Pearson correlation coefficient (r=.84) associated with the dashed regression line is indicated in the top left, in comparison to the solid x=y line; solid x=y line indicates perfect agreement between predicted and measured values. (**e)** Histograms depicting frequency of community richness (number of species in each condition) for experimental designs for DLT0, DTL1, DTL2, and DTL3 (lefthand axis). Circular markers connected by dashed lines indicate the total possible number of combinations for each richness level (righthand axis). Combinations are calculated using the “n choose k” formula with n=15 and k=1 to 15. (**f),(g)** Bar plots showing fraction of conditions in each experimental design cycle containing each species (f) and fiber (g). Species and fibers are sorted left to right along the x-axis (1) alphabetically for DTL0 and by rank order of frequency for DTL1, DTL2, and DTL3. Species bar colors correspond to the legend at bottom right.

For the design of DTL3 and DTL4, we expanded the design space to include discrete, non-equal fiber ratios summing to six parts of 5 g/L fiber in the media (e.g., 3 acacia gum: 2 inulin: 1 xylan). We observed that community compositions of the 15 leave-one-out communities inoculated in each fiber were significantly more similar to one another than to communities cultured in the presence of different fibers (p-value < 1e-10, t-test, **Supplementary Figure 1c,e**). We therefore hypothesized that tuning combinations of fibers with variable ratios could provide a control knob for community composition and function. Including non-equal fiber ratios increased the number of possible conditions in the design space from 2,064,321 to 15,138,354. While DTL2 was designed using an algorithm favoring sets of conditions that maximized both the EIG and the design objective, the expansion of the fiber design space to include varying proportions necessitated the scalability of a Thompson-sampling Bayesian optimization algorithm (**Methods**)^73^. The Thompson-sampling algorithm selected non-equal fiber ratios for 285 of 288 conditions in DTL3. Our experimental results demonstrated that 43 of the top 50 objective conditions contained variable fiber ratios. Therefore, incorporating non-equal fiber ratios can improve the objective compared to using equal parts of each fiber. To determine the reproducibility of community functions, we repeated the top eight objective species-fiber conditions from DTL3. Our results showed consistent results across experimental days and biological replicates (**Supplementary Figure 2**).

To rigorously evaluate improvement in model predictive performance across DTL cycles, we used a consistent test dataset (withheld data used to evaluate prediction performance). Towards this end, we used a cross-validation based approach wherein the model trained only on data from earlier DTL cycles was used to make predictions on test data (data not used to train the model) from all DTL cycles (**Methods**). Including data from subsequent DTL cycles ultimately resulted in improvements in model prediction performance (Pearson correlation between measured and predicted values on held-out data) for all species and metabolites (**Fig. 2c**). Encouraged by the model’s predictive performance across the first two DTL cycles, we expanded the design space beyond the equal fiber proportions used in DTL0,1 and 2 to include all combinations of fiber proportions. After three full DTL cycles, the model’s prediction performance of species abundances achieved a Pearson correlation of at least 0.46, with a median of 0.65 and maximum of 0.79 (p-values < 1e-10) between predicted and measured values (**Fig. 2c**, DTL3) using a standard 20-fold cross validation approach. The Pearson correlation for predicted versus measured butyrate was 0.84 (p-value < 1e-10) (**Fig. 2d**).

To determine if the additional complexity of the MiRNN model was needed to capture the community functions, we performed the same 20-fold cross validation approach as in (**Fig. 2c**) using a linear state-space model with both main-effects (LR) and pairwise-effects (LR2) terms (**Methods**). The MiRNN outperformed the LR for 12 of 15 species and outperformed LR2 for 10 of 15 species and all three metabolites (**Supplementary Figure 3**). Accounting for pairwise-effects (LR2) yielded an improvement in the prediction performance for 11 of the 15 species compared to the LR model. Overall, these results suggest that pairwise-interactions between species and fibers are important for predicting the dynamics of both species and metabolites. The MiRNN, which can capture even more complex, higher-order interactions yielded the best performance.

We evaluated the improvement in prediction performance for the design objective across DTL cycles, where the predicted objective was calculated from model predictions of species abundances and butyrate production (**Methods**). Because model prediction performance of species and metabolites improved over DTL cycles, we observed a corresponding improvement in prediction performance of the design objective across DTL cycles (Pearson correlation = 0.45, 0.70, and 0.77; p-values < 1e-10), with the largest improvement after training on data from DTL2 (**Supplementary Figure 4**). Improvement in the designed objective (**Fig. 2b**) coincided with the substantial improvement in model prediction performance after training on DTL2. We expect that two full DTL cycles were needed to achieve substantial improvement because the model used to design DTL1 was trained only on single-species deletion communities (DLT0) representing a much narrower range of species richness than the full design space.

Species richness is ecological metric to quantify the number of species present in each environment and can trend with certain community functions and increase the degree of functional redundancy^74^. The experimental designs for DTL1 and DTL2 sampled wider ranges of species richness with an emphasis on intermediate community sizes (**Fig. 2e**, blue histograms). The distribution of sampled community richness in DTL1 favored lower richness communities compared to what would be expected from random sampling of the design space (circular markers connected by dashed line, righthand axis) (**Fig. 2e).** In contrast, the distribution of sampled community richness in DTL2 and DTL3 progressively favored higher richness communities, with a mode richness of 10 in DTL3. This shift from low to high richness communities demonstrates the model learned to identify increasingly diverse communities that maintained high butyrate production and compositional persistence across passages.

The frequency that species were selected in each DTL cycle can provide insight into their contributions towards the community functions. Species displayed approximately uniform frequencies in DTL1 and DTL2 (species presence ranging from 28-57% and 40-68% of conditions, respectively) (**Fig. 2f**). However, the most frequent species (*A. caccae*) was present in 100% of conditions in DTL3 whereas the least frequent (*P. copri*) was present in approximately 1% of designs. This trend suggested that the algorithm converged on a subset of species yielding high objective conditions. The frequencies of fiber selection followed a similar trend, with a relatively uniform distribution for DTL1 and DTL2, while DTL3 contained the copresence of inulin and pectin in 99% of conditions and contained acacia gum or arabinogalactan in one condition (0.3% of conditions) (**Fig. 2g**).

Overall, the model achieved accurate prediction performance (Pearson correlation = 0.70, p-value <1e-10) of the design objective after only two full design test learn cycles, demonstrating its ability to map specific initial species-fiber combinations to the compositional, functional, and dynamic properties of synthetic microbial communities (**Supplementary Figure 4**). The data collected throughout the first two DTL cycles consisted of around 700 different species-fiber combinations (around 2000 time-point measurements if accounting for passaging), a relatively small fraction of the 2,000,000 possible species-fiber combinations comprising the entire design space. This result confirms our hypothesis that complex functional and compositional landscapes arising from combinations of dietary fibers and microbial species can be efficiently learned from a comparatively small subset of highly informative samples using a multi-objective Bayesian optimization driven DTL cycle.

### Characterizing the functional landscape of species and fiber combinations

Multi-objective optimization problems typically involve balancing trade-offs between contrasting objectives^75^. For example, a high diversity community may display lower butyrate production due to elevated resource competition than a low diversity community with less resource competition and thus higher butyrate producer growth. To reveal potential trade-offs in our model-guided experiments between butyrate production, Shannon diversity, and temporal dynamics, we evaluated the correlations of each measured objective from all experimental conditions (**Supplementary Figure 5**). While objectives were positively correlated across all comparisons, the highest value of one objective was never the highest in another, suggesting a tradeoff between objective values. Experiments in DTL1 and DTL2 sampled a wide range of butyrate, diversity, and instability values and at times achieved high values of these individual metrics (e.g., high butyrate but low diversity).

In protein engineering, sequence-function landscapes are often used to understand the relationship between the amino acid design space and protein function^76^. To characterize the functional landscape of species-fiber combinations, we encoded species presence-absence, phylogenetic information, and fiber concentrations into a unique vector for each experimentally sampled community and used multidimensional scaling (MDS) to project the set of vectors onto two dimensions (**Methods**). The resulting two-dimensional projection can be visualized as a scatter plot with each community color-coded based on the DTL cycle from which it was sampled or the measurement of the objective value (**Fig. 3a**). The landscape indicates how different species-fiber combinations (represented by location in two dimensions) map to the observed community function (weighted sum of butyrate, diversity, and stability). Large variation in the objective between adjacent points indicates that small changes to the design variables (e.g. presence or absence of individual species) can yield a dramatic change in community function.

**Figure 3.**
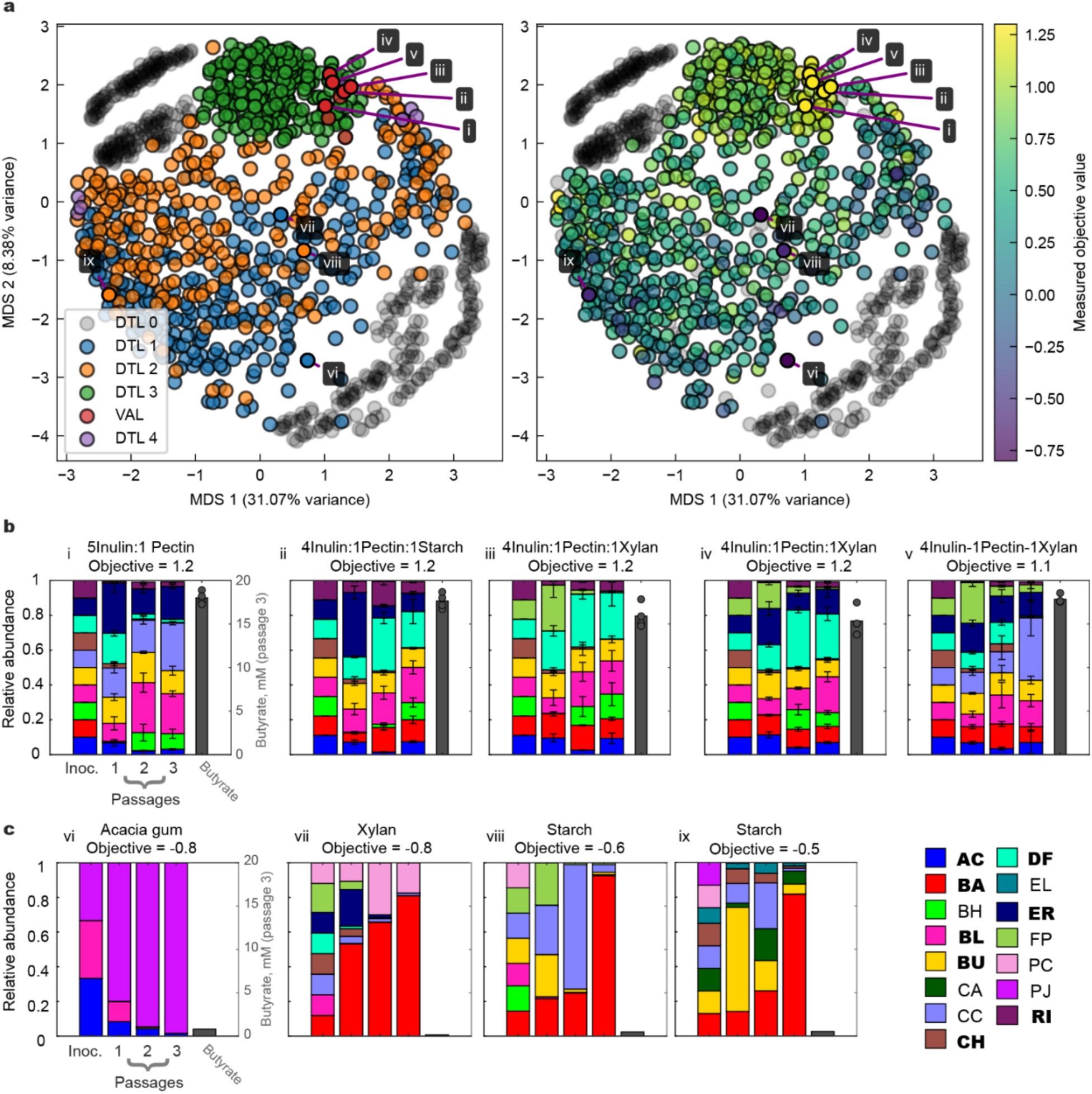
Objective optimization across DTL cycles traverses a rugged functional landscape. (**a)** Objective landscape of DTL cycle and validation data where species-fiber combinations are indicated with gray, blue, orange, green, and red data points for DTL 0, 1, 2, and 3, and DTL3-validation, respectively. The right scatter plot and color bar indicates measured objective values. Markers corresponding to the top five objective bar plots from the validation experiment in panel a are labeled i-v. The axes represent a MDS dimensionality reduction of the species-fiber design space with embedded phylogenetic information (**Methods**). (**b)** Stacked bar plots indicating species composition across passages for the five highest objective communities in the DLT3-validation experiment. Stacked bar height indicates mean relative abundance and error bars indicate standard deviation of n=4 biological replicates. Objective value is indicated as a gray bar, with n=3 biological replicates overlayed as gray circles, corresponding to the righthand axis. Colors indicate species per legend in panel e, bars are stacked alphabetically from bottom to top. Stacked bar plots of relative abundance for each of n=4 biological replicates are shown in Supplementary Figure 2. (**c)** Stacked bar plots indicating species composition across passages for n=1 biological replicates of the four lowest objective communities with formatting matching panel b. Markers corresponding to the lowest objective conditions are labeled vi-ix in panel a.

While much of the landscape exhibited variability in the objective, several top objective communities clustered in a distinct region of the explored design space. To identify clusters in the design space, we performed a k-means clustering algorithm with the number of clusters equal to six, which was selected based on minimizing both the number of clusters and the within-cluster sum of squares (**Fig S6a**). After thresholding the set of conditions to only include those within the 99^th^ percentile measured objective value, at least one condition from two of the original six clusters remained (**Fig S6b**). One cluster shared a common core community of *Anaerostipes caccae, Bifidobacterium adolescentis, Bifidobacterium longum, Bacteroides uniformis, Clostridium hiranonis, Dorea formicigenerans,* and *Roseburia intestinalis*, as well as a common fiber composition of at least four parts inulin and one part pectin (**Supplementary Figure 6c**). By contrast, the other cluster was characterized by presence of *Anaerostipes caccae*, *Blautia hydrogenotrophica*, *Collinsella aerofaciens*, *Prevotella copri*, *Parabacteroides johnsonii*, and starch (**Supplementary Figure 6d**). This result demonstrates that the model discovered at least two different high objective species-fiber combinations and was therefore not limited to sampling a narrow region of the design space. In the first cluster, the top 5 highest objective values were all greater than 1.3. However, in the second cluster, only one condition exceeded an objective of 1.3. This demonstrates that the core species-fiber composition of the first cluster achieved high objective values that were robust to variations in the presence or absence of other species and fibers.

Bayesian optimization algorithms are less susceptible to converging on local rather than global maxima due to the exploration aspect of the acquisition function^77^. While it is difficult to prove that an optimization process has achieved a global maximum in an experimental system, the landscape demonstrates that the algorithm avoided local maxima identified in DTL1 and DTL2 to identify a broader region of high objective values in DTL3 (**Fig. 3a**). Combined with the fact that the model accurately predicted the objective throughout the design space (**Supplementary Figure 4**) and the DTL4 experiments performed following DT3 did not yield substantial improvement (**Fig. 2b**), it is plausible that the region defined by our top objective communities was approaching a global maximum. In sum, Bayesian optimization enabled efficient optimization over a vast, complex functional landscape to identify a robust set of core species-fiber design features that yield high objective values.

### Explainable machine learning offers biological insights

The production of butyrate from dietary fiber is a key function of the healthy gut microbiota^41^. Primary degraders such as Bacteroidetes or Bifidobacterium spp., harbor large genetic arsenals of carbohydrate active enzymes (CAZymes) for utilization of dietary fibers^78,79^. Metabolic byproducts such as acetate and lactate, as well as polysaccharide breakdown products can fuel the growth and metabolic activities of butyrate producers^10^. In addition, certain butyrate producing species can directly utilize specific fibers. For example, *R. intestinalis* has been shown to utilize xylan and many strains of *F. prausnitzii* have been shown to utilize pectin^80,81^. While fiber-mediated interactions are major drivers of human gut community assembly and functions, previous studies have primarily investigated single species or pairwise communities in the presence of single fibers. Analysis of our model trained on 1,185 experimental conditions (27,514 measurements of species and metabolites) could provide deeper insights into influential interactions shaping butyrate production in a more complex multi-species community across different fiber nutrient environments.

Machine learning models such as the MiRNN are often considered “black box” models because of the difficult-to-decipher relationship between inputs and outputs. Shapley additive explanations (SHAP) is an increasingly popular method for quantifying the importance of a particular input variable to model predictions^82^. Specifically, it leverages the computational model to calculate the expected marginal contribution of an input variable to the model prediction of a particular sample condition. Evaluated for all sample conditions (e.g. experimentally evaluated species-fiber combinations), this analysis yields a distribution describing the expected marginal contribution of the input variable on model predictions across all conditions in the dataset.

To gain insight into the mechanisms driving butyrate production, we computed SHAP values to quantify the contribution of each of the fifteen species on butyrate predictions for the experimentally measured conditions (**Fig. 4a**). All known butyrate producers (*A. caccae, C. comes, E. rectale, F. prausnitzii, and R. intestinalis*) have positive median SHAP values, suggesting that all species contributed to butyrate production at the community level. *A. caccae* has the largest median and maximum SHAP values (**Fig. 4a**), suggesting it was the most consistent enhancer of butyrate production (highest mean) and the greatest capacity to promote butyrate production (highest maximum) across many different community-fiber contexts. *A. caccae* is the only species in this community with the capability to produce butyrate from lactate, potentially highlighting the importance of this pathway for butyrate production in complex nutrient environments. This is consistent with a previous study featuring *A. caccae* as the most influential butyrate producer in up to 25-member communities cultured in media containing sugars and lactate^83^.

**Figure 4.**
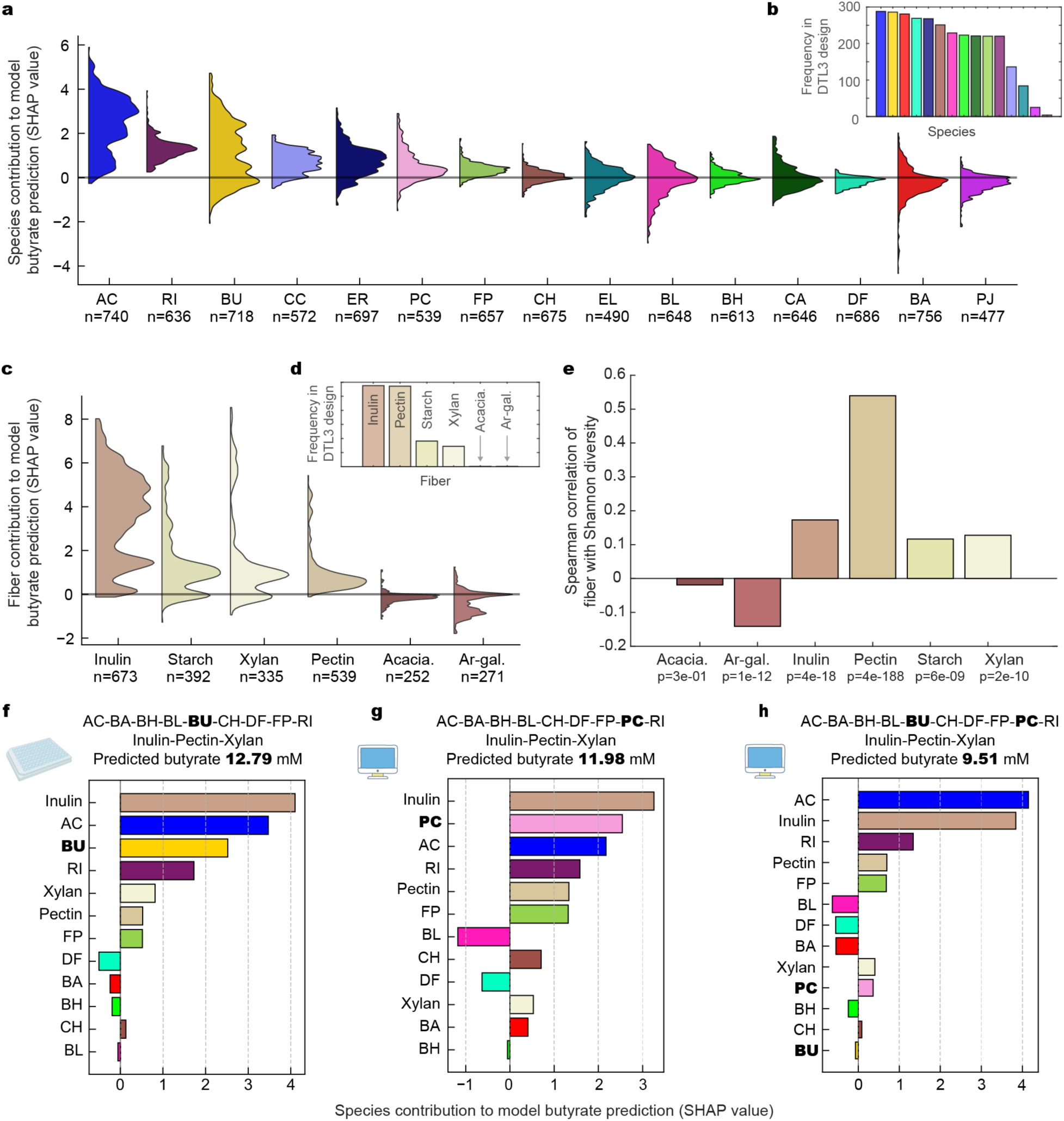
Understanding drivers of butyrate production using explainable machine learning. **(a)** Distributions of SHAP values (Shapley additive explanation) of species contribution to model predicted butyrate. SHAP values are a game-theory based approach to explaining each model input’s contribution to a given model output. Evaluation across training data conditions yields a distribution of values, wherein a broad distribution indicates context-dependent effects of the remaining inputs. SHAP values in the plotted distributions correspond to conditions with the species present, with the number of points in each distribution indicated in the x-axis tick labels. The horizontal line indicates a value of zero. The distributions are sorted by highest to lowest median value. (**b)** Bar plot indicating frequency that each species is represented in the DTL 3 experimental design. (**c)** Distributions of SHAP values (Shapley additive explanations) of fiber contribution to model predicted butyrate for conditions containing respective fiber; formatting matches panel a. Acacia gum and arabinogalactan are abbreviated as “Acacia.” and “Ar-gal.”, respectively. (**d)** Bar plot indicating frequency that each fiber is represented in the DTL 3 experimental design. (**e)** Bar plot where height indicates Spearman rank order correlation coefficient of fibers with Shannon diversity for community data collected throughout DTL cycles; p-values are indicated in x-axis tick labels. (**f)** SHAP explanations of a high objective species-fiber combination (community iii in Fig. 3b) shows high contributions of Inulin, AC, and BU toward butyrate production. (**g)** SHAP explanations of the same community as in f but with BU replaced by PC shows slightly reduced butyrate production where PC has a strong contribution. (**h)** SHAP explanations of the same community with both BU and PC show a drop in predicted butyrate and that the two species have dramatically reduced contributions.

The non-butyrate producing species *B. uniformis* and *P. copri* displayed large SHAP values in comparison to both butyrate and non-butyrate producing species. Previous studies have demonstrated that *B. uniformis* and *P. copri* can metabolize a variety of polysaccharides including inulin and pectin, functioning as primary fiber degraders that can support the growth of butyrate producers^84–87^. In support of this, we found that *P. copri* displayed high growth in the presence of inulin, pectin, and starch single-fiber medium conditions and *B. uniformis* grew well in inulin, pectin, starch, and acacia gum (**Supplementary Figure 1a**). By contrast, out of the five butyrate producers cultured on each of the six single-fiber-containing media, only *A. caccae* and *C. comes* displayed growth in the presence of pectin.

Previous studies have shown certain fibers can promote butyrate production^84^. To evaluate fibers associated with high butyrate production in our data set, we computed the distributions of SHAP values for the contribution of individual inputs to model butyrate predictions (**Fig. 4b**). Inulin, starch, and xylan had relatively large median SHAP values whereas the SHAP values for acacia gum and arabinogalactan were negative. Since SHAP captures statistical relationships between inputs and model predictions, negative values may arise from indirect interactions that reduce butyrate production. Further, SHAP contributions of fibers can be negative due to the fixed initial concentration of fibers, where including one fiber necessarily reduces the initial concentration of other fibers. Consistent with the objective, the four fibers with positive median SHAP values were frequently selected in the DTL3 design, whereas the fibers with negative values were each selected in only one condition (**Fig. 3g**). Pectin was frequently selected in the DTL3 conditions and displayed a relatively low median SHAP value for butyrate production but the second highest median SHAP contribution to predicted diversity behind inulin (**Supplementary Figure 7**). Corroborating this trend, pectin and community Shannon diversity in our experimental data displayed a uniquely large Spearman rank-order correlation, as compared to the other fibers (r > 0.5 and p-value < 1e-10) (**Fig. 4e**).

The requirement for *B. uniformis* in addition to *A. caccae* to yield high butyrate production from inulin has been previously demonstrated in pairwise coculture with inulin as the sole carbohydrate source^27^. The high SHAP contributions of *A. caccae*, *B. uniformis*, and inulin were consistent with the enhancement of butyrate production by *A. caccae* via cross-feeding of polysaccharide breakdown products and/or organic acids as substrates^84^. To further interrogate the context-dependency of these interactions, we analyzed SHAP contributions of a high butyrate producing community (community iii in **Fig. 3b**), which indicated that inulin, *A. caccae*, and *B. uniformis* were the largest contributors to butyrate production (**Fig. 4f**). Interestingly, *P. copri* had a relatively high median SHAP contribution (**Fig. 4a**) but was entirely absent from communities designed in DTL 3 (**Fig. 2f**). When *B. uniformis* was replaced with *P. copri*, the model predicted a moderate reduction in butyrate production, and SHAP analysis attributed inulin, *P. copri*, and *A. caccae* as the largest contributors (**Fig. 4g**). However, when both *B. uniformis* and *P. copri* were included in the community, the model predicted a significantly greater reduction in butyrate production and a substantial reduction in the SHAP contributions of both *B. uniformis* and *P. copri* (**Fig. 4h**). These results suggest that *P. copri* and *B. uniformis* can convert inulin into substrates that support butyrate production by *A. caccae*, but the copresence of both *P. copri* and *B. uniformis* decreases butyrate production and alters their functional roles within the community. This antagonistic interaction impacting butyrate production could be due to a metabolic shift or reduction in species abundance. Corroborating this notion, our results demonstrate that high abundance of *P. copri* (*B. uniformis*) maps to low abundance of *B. uniformis* (*P. copri*) different community contexts (**Supplementary Figure 8**).

To further investigate these interactions, we compared the experimentally measured butyrate concentrations in the subsets of data defined by every combination of presence and absence of *A. caccae*, *B. uniformis*, and inulin (**Fig. 5a**). We first analyzed butyrate data from passage one, because the inoculated species were more likely to be present compared to later passages where certain species may be outcompeted. The butyrate production in the presence of both *B. uniformis*, *A. caccae*, and inulin was significantly higher than other groups (**Fig. 5a**) (one-way ANOVA, p-value < 1e-6, methods). The butyrate measurements in passage three displayed a similar trend of a reduced magnitude (one-way ANOVA p-value < .03) (**Supplementary Figure 9a**).

**Figure 5.**
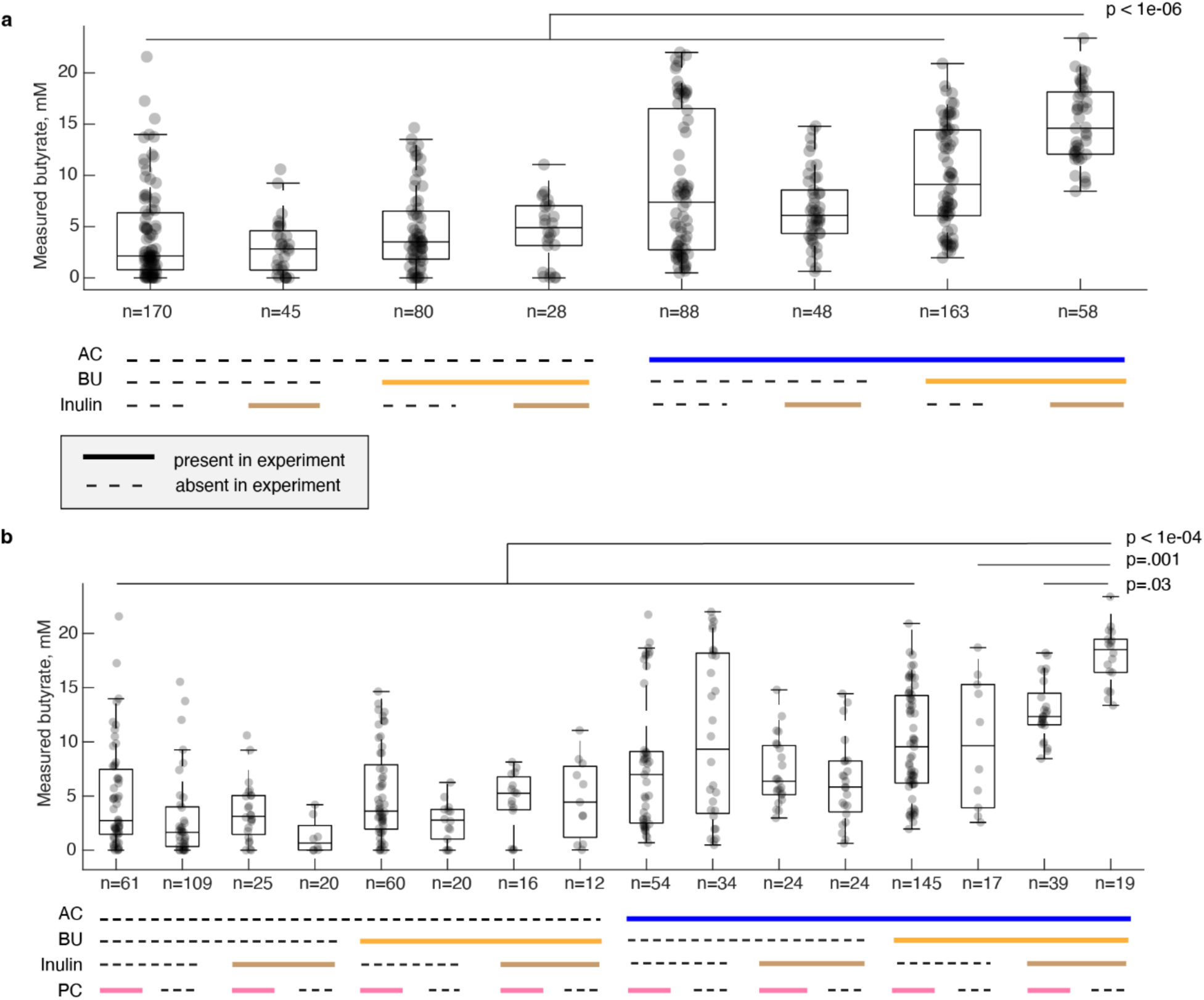
Analysis of experimental butyrate measurements for higher-order interactions. **(a)** Categorical scatter plot of experimentally measured butyrate concentrations for every combination of presence and absence of *A. caccae* (AC), *B. uniformis* (BU), and inulin for passage one. Inulin presence indicates that inulin was the only fiber present in the condition. Presence is indicated by the solid line of which the color corresponds to the variable labeled on the left, and absence is indicated by a dashed black line. Box plots indicate summary statistics of the distribution with center line, box edges, and whiskers indicating median, quartiles, and quartile plus 1.5 times interquartile range, respectively. The statistical comparison at the top of the panel indicates group eight (copresence of AC, BU, and inulin) is significantly higher than all other individual groups using pairwise comparisons (p-values < 1e-06). All pairwise group comparisons are made using one-way analysis of variance (ANOVA) followed by Tukey’s honest significant difference (HSD) procedure using MATLAB’s “anova1” and “multcompare” functions. All 28 pairwise comparisons are reported in **Supplementary Data 4**. (**b)** Categorical scatter plot of experimentally measured butyrate concentrations for every combination of presence and absence of *A. caccae* (AC), *B. uniformis* (BU), inulin, and *P. copri* (PC) for passage one. The statistical comparison at the top of the panel indicates group 16 (copresence of AC, BU, and inulin with the absence of PC) is significantly higher than all other individual groups using pairwise comparisons (p-values ≤ .03). All pairwise group comparisons are made using one-way analysis of variance (ANOVA) followed by Tukey’s honest significant difference (HSD) procedure using MATLAB’s “anova1” and “multcompare” functions. All 120 pairwise comparisons are reported in **Supplementary Data 4.**

To test the hypothesis regarding the antagonistic interactions between *P. copri* and *B. uniformis* on butyrate production, we analyzed the differences in butyrate production for all combinations of presence and absence of the variables. Consistent with the SHAP values, butyrate production is significantly higher in the presence of *A. caccae*, *B. uniformis*, and inulin and absence of *P. copri* than presence of *P. copri* (one-way ANOVA, p-value ≤ .03) (**Fig. 5b**). In addition, there was no significant difference between the presence and absence of *P. copri* for groups that did not contain *A. caccae*, *B. uniformis*, and inulin (one-way ANOVA, p-value > .5) (**Supplementary Data 4**). This trend was not present in passage three (**Supplementary Figure 9b**), consistent with the significantly lower abundance of *P. copri* in passage three than passage one (Mann-Whitney U-test, p-value = .007). In sum, these results demonstrate a higher-order (4^th^ order) interaction involving bacterial species and dietary fiber.

Our results demonstrate that this butyrogenic ecological motif was robust to the myriad of interspecies interactions, species-fiber interactions, and temporal dynamics. *A. caccae* and *B. uniformis* represent the two highest median SHAP contributions to model predicted butyrate of any species and inulin represents the highest median of any fiber. Further, they were the most frequently selected species and fiber in the DTL3 experimental design. By contrast, *P. copri*, which negatively affected butyrate production when combined with *A. caccae*, *B. uniformis*, and inulin, was almost entirely absent from the DTL3 experimental design (**Fig. 2f,g**).

In addition to explaining species and fiber contributions to functions such as butyrate production, we can evaluate the SHAP contributions of inoculated species to predicted abundances of all other species to infer interspecies interactions. To determine the effect of fibers on interspecies interactions, we computed SHAP contributions of inoculated species to predicted growth of species for all experimentally measured conditions and averaged over all conditions that contained a single fiber (**Supplementary Figure 10a**). Of the 210 interactions between species, 137 (65%) changed between positive and negative values across environments with different individual fibers, demonstrating a fiber-dependent inter-species interaction network (**Supplementary Figure 10b**), consistent with a previous study that investigated how single fibers shaped the inter-species interaction network^88^. In particular, *Eggerthella Lenta* had a positive SHAP contribution to the growth of BU that was robust across all 6 fibers, consistent with its positive impact on *Bacteroides spp.* in other nutrient environments and community contexts^26^. Consistent with the complex interactions between *B. uniformis*, *P. copri* and *A. caccae* in the presence of inulin, *B. uniformis* and *P. copri* displayed positive SHAP contributions on *A. caccae*. Further, *B. uniformis* and *P. copri* exhibited bidirectional negative SHAP contributions in the presence of inulin. In sum, explainable machine learning revealed higher-order interactions and uncovered *A. caccae*, *B. uniformis*, and inulin as a robust motif for butyrate production across fiber environments.

### Investigating colonization and metabolite profiles of designed species-fiber combinations in a gnotobiotic mouse model

While holding tremendous therapeutic potential, the colonization and impact of defined bacterial therapeutics is unpredictable, as evidenced by recent clinical trials^89,90^. The administration of dietary fibers as prebiotics is a promising approach for modulating the human gut microbiome but also suffers from unpredictable outcomes^8^. To enhance predictability of outcomes, co-administration of defined consortia and specific microbial accessible carbohydrates have the potential to provide a metabolic niche and thus enhance engraftment and target functions in the human gut^91^.

To investigate the effects of the model-designed species-fiber combinations, we orally gavaged gnotobiotic mice with specific communities and fed mice tailored defined diets containing the model-designed dietary fibers. Using this approach, we characterized the temporal changes in community composition and functions of interest of specific model-optimized species-fiber combinations in the mammalian gut (**Fig. 6a**). We selected a high versus low design objective species-fiber combination to determine potential differences in their properties within the mammalian gut. For the high objective condition, we selected the community consisting of *A. caccae, B. adolescentis, B. hydrogenotrophica, B. longum, B. uniformis, C. hiranonis, D. formicigenerans, F. prausnitzii,* and *R. intestinalis* and fiber ratio of 4 inulin: 1 pectin: 1 xylan (HiComm-IPX) (**Fig. 3b** iii). HiComm-IPX contains a core set of species and fibers that resulted in measured objective values that were robust to changes in other species and fibers (**Supplementary Figure 7**). The low objective community consisted of *B. adolescentis, B. longum, C. comes, C. hiranonis, D. formicigenerans, E. rectale, F. prausnitzii,* and *P. copri* and xylan as the fiber (LoComm-XYL). This community included a similar number of total species to HiComm-IPX, a similar number of butyrate producers, and did not contain starch. Since starch can be degraded by host enzymes, the dynamics and functions of the community may be less predictable due to host-fiber interactions as opposed to the microbial accessible carbohydrates present in the high-objective condition (**Fig. 3c** vii). To determine the effects of the model-optimized fiber profile on community properties, we orally gavaged a third group of mice with the HiComm and the mice were fed a defined diet with cellulose (HiComm-CEL). Cellulose is not accessible to human gut bacterial species and thus represents the absence of microbial accessible carbohydrates (i.e. no fiber)^92^. Using fecal samples, we performed time-series measurements of community composition over a 15-day study to reveal community dynamics. Community composition was determined via 16S rRNA gene sequencing. In addition, we measured volatile fatty acids as well as the total colony forming units (CFUs) to assess the absolute abundance of viable cells as a metric for colonization (**Methods**).

**Figure 6.**
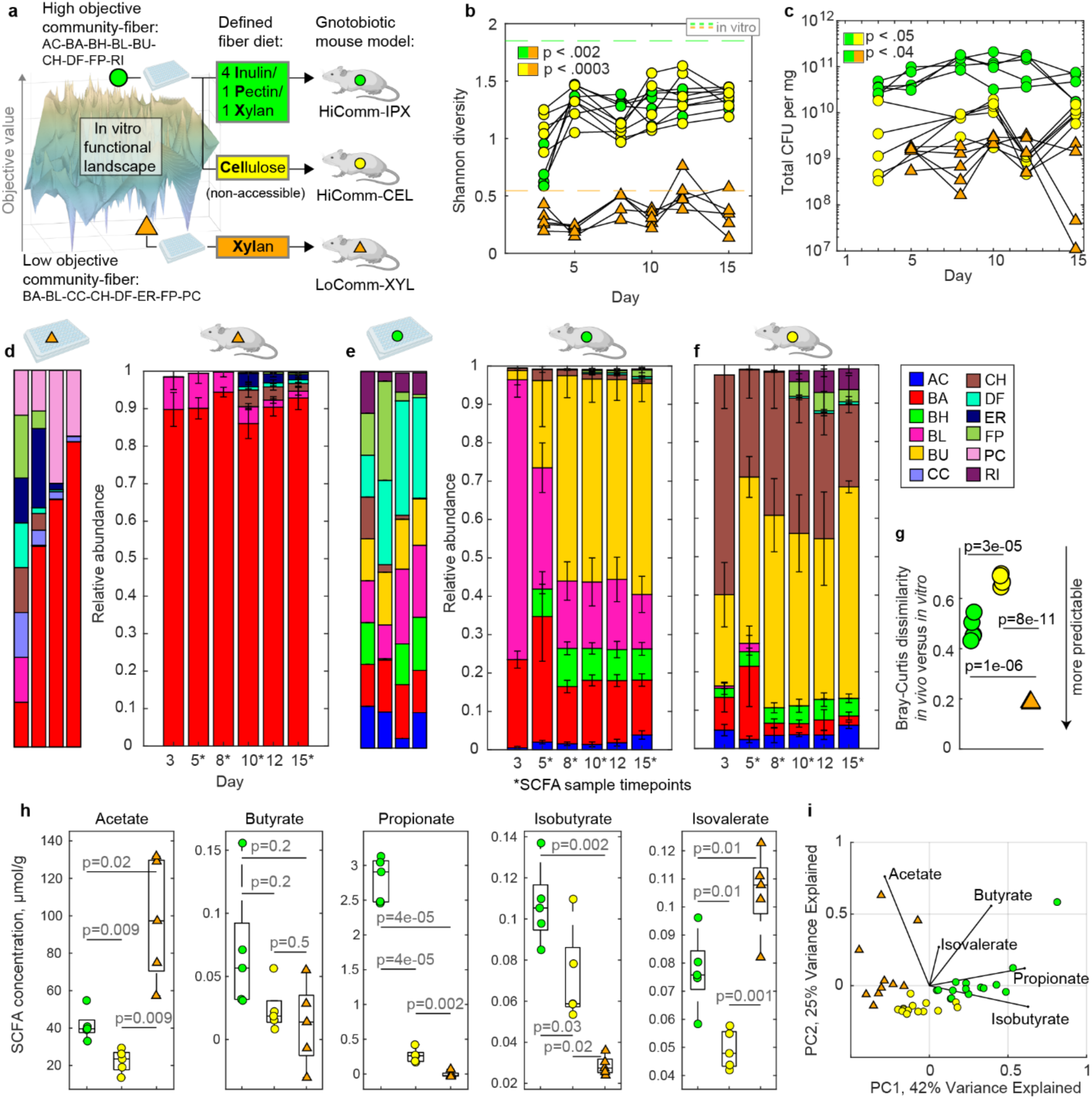
Composition, function, and dynamics of designed species-fiber combinations in a gnotobiotic mouse model with defined fiber diets. **(a)** Study design in which germ-free mice are colonized with a high design objective community with and without its corresponding designed fiber proportions in defined diets, as well as a low objective community with corresponding fiber diet (Methods). Groups of n=5 mice were gavaged with even strain ratios on days one and eight of the 15-day study. Fecal samples were collected every two to three days. **(b)** Line plot indicating *in vivo* community Shannon diversity over time for n= 5 biological replicates. Marker style indicates community and marker color indicates diet according to panel a. Horizontal dashed lines indicate *in vitro* passage three diversity of corresponding communities. The range of p-values for unpaired, two sample t-tests with Benjamini-Hochberg multiple comparison adjustment is indicated if significant for all timepoints; all p-values are shown in **Supplementary Table 2**. **c** Line plot indicating viable population density (colony forming units, CFU/mg, limit of detection is 10^5^) over time for n=5 biological replicates per group (Methods). The range of p-values for Welch’s t-test with Benjamini-Hochberg multiple comparison adjustment is indicated if significant for all timepoints; all p-values are shown in **Supplementary Table 2**. Marker style and colors indicate groups according to the legend in panel a. **(d)-(f)** Stacked bar plots of community composition where height indicates mean relative abundance of species and error bars indicate standard deviation over n=5 biological replicates. All replicates are shown in **Supplementary Figure 11a**. Lefthand stacked bar plots show corresponding *in vitro* relative abundance over passages of communities as detailed in Fig. 4c iii and Fig. 4e vii. Colors indicate species according to legend in the lower righthand portion of panel f. **(g)** Categorical scatter plot indicating Bray-Curtis dissimilarity between day 15 community composition for in *vivo* communities and final passage of corresponding *in vitro* community. The p-value for an unpaired, two sample t-test between groups of n=5 replicates with Benjamini-Hochberg adjustment for multiple comparisons is shown above the plot. **(h)** Categorical scatter plots of fecal short chain fatty acid (SCFA). Markers indicate the average concentration of SCFA for a given mouse over available samples from days 8-15 corresponding to when community composition and metabolites had converged to approximately steady state. All available SCFA data is shown in **Supplementary Figure 11b**. Box plots indicate summary statistics for n=5 replicates where middle line indicates median, box edges indicate quartiles, and whiskers indicate maximum and minimum datapoints. The p-values for Welch’s t-test adjusted for multiple comparisons via Benjamini-Hochberg procedure are indicated in the plot. Marker style and color is indicated in panel a legend. **i** Scatter plot indicating vector weightings (lines) and data scores (markers) for first two components of a principal component analysis (PCA) on approximately steady state SCFA concentrations (available samples from days 8-15). Horizontal and vertical axis labels indicate percent variance described by corresponding component; marker style and color is according to panel a legend.

We denote the species-fiber conditions as e.g. HiComm-IPXc or HiComm-IPXm corresponding to *in vitro* culture or mouse, respectively. Our results demonstrated that HiComm-IPXm and HiComm-CELm displayed significantly higher Shannon diversity than LoComm-XYLm across all time points, consistent with the *in vitro* experimental trends (unpaired two-sample t-test adjusted for multiple comparisons, p-values < .002, **Supplementary Table 2**) (**Fig. 6b**). HiComm-IPXm also displayed significantly higher colonization of viable bacteria (CFU/mg) than LoComm-XYLm across all time points (unpaired two-sample t-test adjusted for multiple comparisons, p-values < .04, **Supplementary Table 2**) (**Fig. 6c**). Although HiComm-IPXm and HiComm-CELm did not significantly differ in Shannon diversity across the timeseries, the total CFU/mg revealed significantly higher colonization of viable bacteria for the HiComm-IPXm than for HiComm-CELm (**Fig. 6c**, **Supplementary Table 2**, adjusted p-values < .05). Therefore, the optimized species-fiber combination was necessary to simultaneously achieve both high Shannon diversity and a high degree of mammalian gut colonization by bacteria. These properties could enhance the health-beneficial effects of probiotic consortia^21,93^.

Our results demonstrated substantial differences in species composition between HiComm-IPXm and HiComm-CELm, indicating a strong impact of the designed fiber diet on community composition and species engraftment (**Fig. 6e,f**). To quantify whether the designed fiber diet in HiComm-IPXm resulted in species composition that was more similar to its *in vitro* counterpart than HiComm-CELm and thus more predictable, we computed a Bray-Curtis dissimilarity metric between the in *vivo* compositions at the final timepoint and *in vitro* composition at the final passage (**Fig. 6g**). HiComm-IPXm was significantly more similar to its *in vitro* counterpart than HiComm-CELm (p-value = 3e-05, unpaired two-sample t-test adjusted for multiple comparisons). For example, *B. longum* represented about 10% relative abundance in both HiComm-IPXc and HiComm-IPXm but was effectively absent from HiComm-CELm (**Fig. 6e,f**). Similarly, *C. hiranonis* was low relative abundance in HiComm-IPXc and in HiComm-IPXm, while it represented greater than 20% relative abundance across the HiComm-CELm timeseries. LoComm-XYLm was also significantly more similar to its *in vitro* counterpart than HiComm-CELm (t-test, p-value = 8e-11). Mirroring *in vitro* data, *B. adolescentis* converged to a high relative abundance in the LoComm-XYLm group, while this species displayed modest relative abundance in HiComm-IPXm (**Fig. 6d**). In sum, these results demonstrate certain consistent compositional trends *in vitro* and *in vivo* and highlight that fiber can improve the predictability of fiber-species microbiome interventions.

To understand the metabolite profiles of our designed species-fiber combinations, we measured the fecal short chain fatty acid (SCFA) profile, which included acetate, butyrate, propionate, isobutyrate, and isovalerate over time. We analyzed SCFA profiles corresponding to available timepoints for which the species composition had stabilized for all groups (days 8, 10, and 15). We found substantial variation with community and diet, indicating that designed combinations of species and fibers shaped metabolic function *in vivo* (**Fig. 6h**). The concentrations of SCFAs were higher in HiComm-IPXm than HiComm-CELm, but this difference was not statistically significant for butyrate (p-values < 0.03 and equal to 0.2, respectively, Welch’s t-test adjusted for multiple comparisons with Benjamini-Hochberg procedure). Butyrate concentrations were very low overall, resulting in noisy measurements. In addition, since butyrate is a primary energy source for colonocytes, the measured butyrate is determined by production by the gut community and host uptake and thus may not accurately reflect the butyrate production capability of the community^21^. HiComm-IPXm displayed higher in butyrate, propionate, and isobutyrate (Welch’s t-test, p=0.2, 4e-5, and 0.002), whereas LoComm-XYLm exhibited significantly higher acetate and isovalerate (Welch’s t-test, p=0.009 and 0.001). A principal component analysis (PCA) on the SCFA data from days 8 to 15 identified two components that account for around 70% of the total variance (**Fig. 6i**). Data from each group clustered together in the PCA plot, reflecting differences in their functional profiles.

Overall, the SCFA profile of HiComm-IPXm exhibited elevated levels of multiple health beneficial metabolites. For instance, isobutyrate produced by gut microbiota has been associated with positive outcomes in ulcerative colitis treatments in human subjects^94^. Further, isobutyrate has anti-inflammatory properties and is a preferred energy source for colonocytes, similar to butyrate^95,96^. Microbiota derived propionate has a range of health beneficial properties including regulation of gluconeogenesis, protection against atherosclerosis, and reduction of weight gain^97–99^. By contrast, LoComm-XYLm displayed high levels of acetate (reaching almost 300 umol/g on day 8) as well as significantly higher levels of isovalerate (**Supplementary Figure 11b)**. Acetate plays a complex and context-dependent role in human health, being linked to both beneficial and detrimental effects across a range of biological functions^100^. High levels of acetate have been associated with hepatic lipogenesis, while isovalerate has been associated with depression^101^. The observed high levels of acetate and isovalerate in LoComm-XYLm were likely a result of the high proportion of *B. adolescentis* (*Bifidobacteria* are prominent acetate producers) and low ecological diversity^18^. Overall, the high abundance of *B. adolescentis* and thus low ecological diversity in LoComm-XYLm may lead to over or underproduction of key health-relevant metabolites produced by a more diverse community. In sum, the diversity and composition of designed species-fiber combinations displayed similar trends *in vitro* and *in vivo* and the treatments yielded substantially different SCFA profiles and thus community metabolic states.

## DISCUSSION

The integration of high-throughput experiments, computational modeling, and advanced optimization techniques holds great promise for guiding precision microbiome interventions to achieve desired functional states. We applied a Bayesian optimization framework to navigate the space of fiber concentrations and initial species abundance to design complex synthetic communities with desired functional outputs. Our results that fiber selection is a critical control point for tailoring community functions. Our model accurately predicted butyrate production across various fiber and community combinations over a series of three 48-hour passages. Our model also achieved strong prediction performance of the objective function after training on around 1,000 species fiber combinations over three passages (around 1800 measurements). Our experimental dataset represents a small percentage (0.008%) of the over 15,000,000 possible species-fiber combinations, demonstrating that the rules governing community assembly and function can be learned from a relatively small number of highly informative conditions using a Bayesian optimization driven design-test-learn cycle. Further, community composition and diversity of species measured *in vitro* were consistent with our *in vivo* results.

While higher order interactions shaping microbial community functions can play influential roles in system properties, there are limited methods for identifying these interactions from data^102,103^. The presence of higher order interactions highlights the importance of using a flexible model that can capture complex interactions involving both species and resources, in contrast to commonly implemented ecological models such as generalized Lotka-Voltera (gLV) or MacArthur’s consumer resource (MCR)^26,104,105^. The ability of the MiRNN to outperform a linear model in prediction of both species and metabolites, even when accounting for pairwise interactions, demonstrates the presence of higher-order interactions that govern the relationship between species, metabolites, and fibers (**Supplementary Figure 3**). Further, accounting for all pairwise-interactions between species and fibers resulted in a model (LR2) with 6,194 parameters, which was more than three times the number of parameters of the MiRNN, which had 2,515 parameters. Therefore, when predicting multiple output variables, neural networks do not necessarily have more parameters than seemingly simpler linear models, which suffer from the curse of dimensionality when accounting for interactions. By combining flexible models such as neural networks with explainable machine learning approaches, we can elucidate complex rules governing the functional landscape of butyrate production involving higher order interactions between species and fibers.

Ecological insights from our model and data reveal that the widely referenced paradigm of synergistic butyrate production from fiber, facilitated by cross-feeding between primary degraders and butyrate producers, plays a robust and influential role in complex communities^21^. In support of this paradigm, fiber degrading species such as *B. uniformis* and *P. copri* displayed growth in the presence of inulin in monoculture and had positive SHAP contributions to the growth of several butyrate producing species including *A. caccae*. However, not all fiber degrading species promoted the growth of butyrate producers. For example, while *B. longum* exhibited growth in the presence of inulin in monoculture, *B. longum* had negative SHAP contributions to the growth of all butyrate producing species (**Supplementary Figure 1, 10**). The quantification and statistical validation of ecological motifs involving *A. caccae*, *B. uniformis*, *P. copri*, and inulin demonstrates that fourth-order interactions between species and fibers are robust to community context and are key drivers of complex community properties. The substantial decrease in the SHAP contribution of *B. uniformis* and *P. copri* on butyrate production in communities with both species may arise from an interaction-driven metabolic shift that reduces the ability of *A. caccae* to produce butyrate. The ratio of *Bacteroides* and *Prevotella* has drawn recent attention as a microbiome biomarker indicative of diet, lifestyle, and disease state^106,107^. Our results suggest that the *Bacteroides* to *Prevotella* ratio affects butyrate metabolism *in vitro*. Elucidating the mechanisms of these interactions remains a future area of research.

While our results demonstrate that the incorporation of fibers improved the similarity between *in vitro* and *in vivo* community compositions, predicting *in vivo* species engraftment from *in vitro* data remains a challenge. For example, while the butyrate producing species *F. prausnitzii* was present in each passage of the *in vitro* HiComm-IPXc condition, *F. prausnitzii* only colonized certain mice following a second gavage (day 8) in HiComm-IPXm (**Fig. 6e,f**). This suggests that the ability of *F. prausnitzii* to colonize depends on modifications of the murine gut environment by the community, consistent with a previous study^108^. An avenue of future research is to “close the loop” between *in vitro* and *in vivo* design-test-learn cycles by incorporating information from animal models for further refinement of community design goals. Further, while we relied exclusively on experimental data to train the MiRNN, the incorporation of prior knowledge about species consumption of fibers or production of metabolites could be leveraged to improve initial model predictions. Integrating high-throughput *in vitro* data, data from *in vivo* animal models, and prior knowledge is a plausible extension of the framework presented in this study that could improve our ability to predict and control the microbiome and influence host phenotypes across different environmental contexts.

In sum, we establish a blueprint for the bottom-up design and optimization of function, composition, and dynamics in defined microbial communities using both species and resources as influential control points. Further, the ability to translate community composition and functional properties designed *in vitro* to the mammalian gut microbiome constitutes an important step towards the bottom up, rational design of functional species-fiber combinations (synbiotics) to benefit human health. Because our machine learning-based approach is flexible to any species and fiber inputs, as well as any measurable community property as an optimization target, it could readily be implemented as a pipeline for high-throughput design and optimization of candidate synbiotic formulations for evaluation in preclinical animal models. The ability to optimize microbial communities by tuning specific dietary fibers could provide a critical advance towards realizing the true potential of living bacterial therapeutics. Finally, this broader approach of designing and optimizing microbial communities and key environmental factors to promote their colonization or target functions could address challenges in agriculture, biomanufacturing, and wastewater treatment over the coming decades.

## METHODS

### Strain, media, and growth conditions

The following methods are adapted from Hromada 2021, Clark 2021 and Venturelli 2018.^109–111^ All anaerobic culturing was carried out in a custom anaerobic chamber (Coy Laboratory Products, Inc) with an atmosphere of 2.5 ± 0.5% H2, 15 ± 1% CO2 and balance N2. All prepared media, stock solutions, and materials were placed in the chamber at least overnight before use to equilibrate with the chamber atmosphere. The species used in this work were obtained from the sources listed in **Supplementary Data 2** and permanent stocks of each were stored in 25% glycerol at −80 °C. Batches of single-use glycerol stocks were produced for each species by first growing a culture from the permanent stock in anaerobic basal broth (ABB) media (HiMedia or Oxoid) to stationary phase, mixing the culture in an equal volume of 50% glycerol, and aliquoting 400 μL into Matrix Tubes (ThermoFisher) for storage at −80 °C. Quality control for each batch of single-use glycerol stocks included (1) plating a sample of the aliquoted mixture onto LB media (Sigma-Aldrich) for incubation at 37°C in ambient air to detect aerobic contaminants and (2) next-generation DNA sequencing of 16S rDNA isolated from pellets of the aliquoted mixture to verify the identity of the organism (Illumina).

For each experiment, precultures of each species were prepared by thawing a single-use glycerol stock and combining the inoculation volume and media listed in **Supplementary Data 1** to a total volume of 5 mL for stationary incubation at 37°C. Incubation times are listed in **Supplementary Data 2**. Precultures were sub-cultured by inoculating 5 mL of fresh media with cultures grown for the specified growth time at an OD600 target of 0.01. Subcultures were grown for 24 hours then used to inoculate community and monoculture experiments. Subculturing of precultures was performed to reduce variability of growth phase for inocula for community experiments.

### Gnotobiotic mouse experiments

All germ-free mouse experiments were performed following protocols approved by the University of Wisconsin-Madison Animal Care and Use Committee. Male C57BL/6 gnotobiotic mice (wild-type) of age 9-11 weeks were fed the defined diets based on *in vitro* experimental results. Three defined diets were used in this experiment: Inulin-pectin-xylan fiber diet (Envigo, TD.240103), xylan diet (Envigo, TD.240104, cellulose diet (Envigo, TD.240180) (**Supplementary Data 3**). Mice were transferred from isolaters to biocontainment cages (Allentown Inc.) and started feeding with defined diets five days prior to receiving community gavage. Transfer and all subsequent handling of mice was performed in biosafety cabinets.

Mice received 200 uL of bacterial inoculum normalized to 0.125 OD600 via oral gavage. Inoculum contained equal species proportions by optical density in 1x Phosphate buffered saline (OD600 of each species in the mixure was targed at 0.125/n where n indicates number of species in the respective community. Identical gavages were performed on Day 1 and Day 7 of the study. Inocula mixtures for gavage were transferred to Hungate tubes (Chemglass) on ice; remaining mixture was sampled for DNA sequencing and plated aerobically to check for aerobic contaminants. for the duration of the experiment. Groups of n=5 mice corresponding to a given diet/community were co-housed in a single cage. Fecal samples were collected every 2-3 days after oral gavage and transported on ice. At the end of the experiment, mice were euthanized, and the cecal contents were collected for NGS sequencing and CFU plating.

### The Microbiome Recurrent Neural Network (MiRNN)

To model the dynamics of species abundance and metabolite concentrations over passages, we used the Microbiome Recurrent Neural Network (MiRNN), a machine learning model tailored to predict microbial community dynamics that eliminates the possibility of predicting physically unrealistic species abundances and metabolite concentrations. The MiRNN can be trained to predict changes in species abundance and metabolite concentrations over time in response to inoculation densities and selection of fibers. Once trained, the MiRNN can be used to select experimental conditions predicted to maximize butyrate production and community diversity and minimize the variation in species abundances over passages.

The MiRNN is a recurrent neural network that uses a constraint to ensure that species that were absent at previous time steps remain absent from the community. The model equations are given *by*,

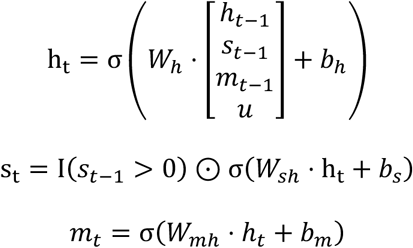

where 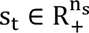 is a vector of species abundances, 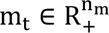 is a vector of metabolite concentrations, 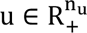 is a vector of fiber concentrations, 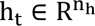 is a vector of latent variables that inform predictions of species and metabolite concentrations at subsequent time steps, and σ is the rectified linear unit activation function. The set of neural network parameters is denoted as θ = {W_h_, b_h_, W_sh_, b_s_, W_mh_, b_m_, h_0_} ∈ R^nθ^. The indicator function I(s_t−1_ > 0) forces predictions of species abundances at time *t* to be zero if the species abundances at time *t* − 1 were zero.

To enable a more compact notation, we denote the input to the MiRNN as q = (s_0_, m_0_, u, τ), which specifies the initial species abundances, metabolite concentrations, fibers, and passage number, τ. The MiRNN’s predictions of species abundances and metabolite concentrations are denoted as 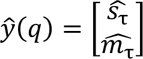 Measurements of species abundances and metabolite concentrations corresponding to the input *q* are denoted as 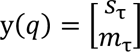.

### Linear state-space models (LR and LR2)

Linear state-space models predict species and metabolites in the next time step as a linear combination of their values in the previous time step. The linear state-space model with main effects (LR) is,

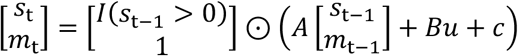

where 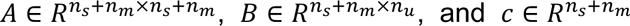. Like in the MiRNN, the indicator function I(s_t−1_ > 0) forces predictions of species abundances at time *t* to be zero if the species abundances at time *t* − 1 were zero. To account for interactions between species, metabolites, and fibers, we can introduce the following basis function,

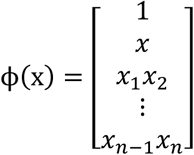

Using this basis gives a linear state-space model with both main effects and pairwise effects (LR2),

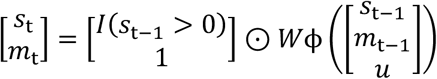

where *W* ∈ *R*^*n_s_*+*n_m_*×*n_b_*^. The dimension of the basis is 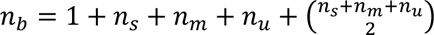. In this study, with 15 species, 3 metabolites, pH (considered a metabolite), and 6 fibers, the dimension of the basis is 326. Because there are 19 outputs including species, metabolites, and pH, the LR2 model has 6,194 parameters (dimension of the weight matrix *W*).

### Parameter estimation

To estimate model parameters, θ, we use an approximate Bayesian inference method to infer a parameter distribution by conditioning on experimentally observed data. We define a set of *n* experimental conditions as Q = {*q*_1_,…, *q*_*n*_} and a set of corresponding measurements of species and metabolite measurements as *D*(*Q*) = {*y*(*q*_1_),…, *y*(*q*_*n*_)}. We use the Laplace approximation to determine the mean and covariance of a Gaussian distribution that approximates the true parameter posterior. A prior parameter distribution centered at zero is used to promote model simplicity, denoted as p(θ) = *N*(0, α^−1^I). The precision of the prior, α, is a hyper-parameter that determines how much parameters are penalized for deviating from zero. Because measurements of species and metabolites are affected by many potential sources of noise, we use the model to estimate the expected value of species abundances and metabolite concentrations and assume that the distribution due to measurement noise is Gaussian. We therefore estimate the distribution of species and metabolites conditioned on a particular experimental condition *q* to be 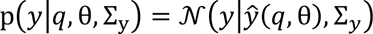. The covariance matrix Σ_*y*_ is a diagonal matrix where 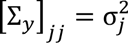 is the variance of the measurement noise associated with species or metabolite *j*, where j = 1,…, n_s_ + n_m_. Given this model for the distribution of species and metabolites, the likelihood of observing a particular set of measured values *D*(*Q*) is

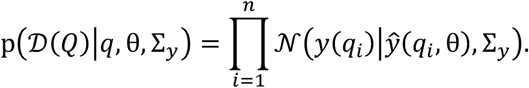

Bayes theorem defines the posterior parameter distribution as proportional to the likelihood of a dataset multiplied by the parameter prior distribution,

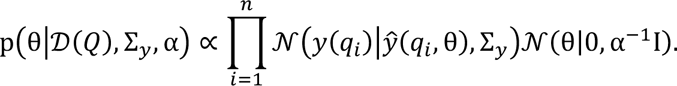

Maximizing the log of the parameter posterior distribution with respect to θ gives the *Maximum A Posteriori* (MAP) estimate,

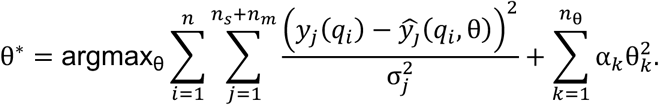

The Laplace method approximates the posterior parameter distribution as a Gaussian with a mean of θ^∗^ and covariance as the inverse of the matrix of second derivatives of the negative log posterior,

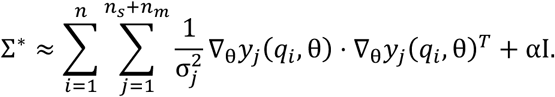

With the posterior parameter distribution approximated as a Gaussian, p(θ|*D*(*Q*), Σ_*y*_, α) = *N*(θ|θ^∗^, Σ^∗^), the posterior predictive distribution is determined by marginalizing with respect to the posterior,

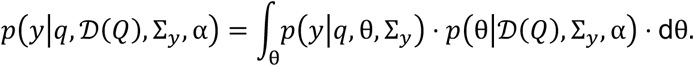

We use the Expectation-Maximization (EM) algorithm to optimize the precision of the parameter prior, α, and the measurement precision, Σ_*y*_, which collectively control model regularization by weighting deviations from the prior and likelihood, respectively^61^.

### Cross-validation to evaluate model prediction performance

To evaluate the accuracy of the MiRNN as data is collected in each DTL cycle, we used a two-step cross-validation approach. When evaluating prediction performance of the model using data up to a particular DTL cycle, we first perform 20-fold cross-validation on all data collected up to that DTL cycle. In the second step, we train the model on all data up to that DTL cycle and use data from all subsequent DTL cycles as a test set. Combining the held-out predictions from cross-validation and the test predictions from subsequent DTL cycles provides model predictions for all samples in the entire data set. By comparing model predictions to measured values, we can evaluate how much incorporating data from subsequent DTL cycles improves model prediction performance in a way that ensures the performance metric is consistent across all comparisons.

### Bayesian optimization

Once the MiRNN is trained on previously collected data, we can use the model to predict butyrate production, community diversity, and stability in previously untested conditions. We define an objective function, J(*q*, θ^∗^), which quantifies a weighted combination of predicted butyrate production, stability, and community diversity,

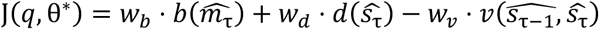

where 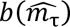 is the predicted butyrate at passage τ, 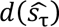 is the Shannon diversity of the predicted community at passage τ, and 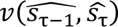 computes instability as the average standard deviation in species abundances between passages τ − 1 and τ. The coefficients *w*_*b*_, *w*_*d*_, and *w*_*v*_ weight each component of the objective function based on the highest observed values so that each objective is given equal priority in the combined objective function. Bayesian optimization combines the design objective J(*q*, θ^∗^) with a measure of the information content of a set of experimental conditions to explore undersampled regions of the design space.

### Bayesian optimization using the expected information gain

We used an experimental design strategy called Bayesian optimization that seeks a set of experimental conditions that accomplish the dual task of improving the accuracy of the model while also improving the design objective. One way to quantify how well a data set will help improve a model is to use the expected information gain (EIG), which quantifies how much a data set is expected to constrain model parameter estimates according to the determinant of the expected parameter covariance matrix. For a set of experimental conditions, *Q* = {*q*_1_,…, *q*_*n*_}, the EIG can be approximated as,

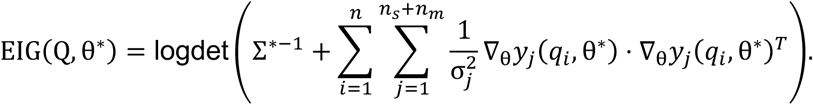

which quantifies how much the data is expected to minimize the volume of the posterior parameter distribution. We used a Greedy algorithm to optimize experimental designs that maximize a combination of the design objective and the EIG, where experimental condition *q*_1_ is the condition that maximizes J(*q*, θ^∗^) and all subsequent conditions *q*_*k*_ maximize

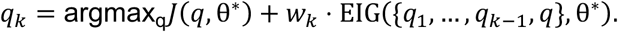

The weighting coefficient *w*_*k*_ is gradually increased from an initially small value (1 × 10^−3^) until a unique condition is selected, *q*_*k*_ ∉ {*q*_1_,…, *q*_*k*−1_}. To select experimental conditions for DTL 1, we used the above Bayesian optimization algorithm without the design objective to focus entirely on exploration (maximization of the EIG) and incorporated the design objective to select experimental conditions in DTL 2.

### Bayesian optimization using Thompson sampling

In DTL 3, we expanded the design space to include unequal fiber ratios (e.g. 3 parts inulin, 2 parts pectin, and 1 part starch instead of equal fractions of each fiber), which increased the design space from 2,064,321 possible experimental conditions to 15,138,354. Due to the computational cost of computing the EIG, we used a Bayesian optimization algorithm called Thompson sampling to select experimental conditions in DTL 3 as a more scalable alternative^73^. Thompson sampling exploits and explores an objective function by first sampling candidate parameter values from the posterior parameter distribution, θ^(k)^ ∼ *N*(θ^∗^, Σ^∗^), and then uses the resulting model to identify the condition that maximizes the design objective. For each parameter sample, *k* = 1,…, *n*, an experimental condition is selected by optimizing the design objective

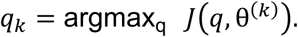

The process of sampling and selecting a condition was repeated until *n* = 288 unique conditions were selected (corresponding to three 96 well plates).

### Synthetic community experiments

All anaerobic culturing was carried out in a custom anaerobic chamber (Coy Laboratory Products, Inc) with an atmosphere of 2.5 ± 0.5% H2, 15 ± 1% CO2 and balance N2. All prepared media, stock solutions, and materials were placed in the chamber at least overnight before use to equilibrate with the chamber atmosphere.

All media contained 5 g/L of total fiber. Concentrated fiber stock solutions were prepared at 15.2 g/L (∼3x working concentrations) as follows: a paste was made by mixing the desired mass of fiber with around 15% of the final target volume of water, using a stir bar to mix as thoroughly as possible. Hot water was added to the target volume while mixing continued. The fibers were autoclaved for 30 minutes at 121 degrees Celsius in bottles containing the stir bars. Upon completion of the autoclave cycle, the fibers were immediately stirred each using the stir plate to fully dissolve the fibers. Duration of stirring was about five minutes, except for the starch which was stirred for between 1 and 16 hours. One day prior to inoculation, the fiber combinations were assembled in deep well blocks using a Tecan Evo liquid handling robot.

Community cultures were performed in 2.2 mL deep well blocks at 1200 uL scale. Base medium lacking carbohydrates was prepared at 1.75x concentration and 685 uL was added to the 395 uL of concentrated fiber mixture in the deep well blocks. The inoculation density of each species was held constant across all conditions containing that species. The constant inoculation density was calculated based on inoculating the full 15-member community at a total target OD600 of 0.01, or 0.01/15 = 0.00067 OD600. In other words, the total inoculation density was proportional to the number of species in the condition, with the total inoculation density of monocultures at 0.00067 OD600 and the full 15-member community at 0.01 OD600. Sub-cultured precultures prepared as previously described were normalized to 0.1 OD600 in 1x base media lacking carbohydrates. A Tecan Evo liquid handling robot was used to assemble communities according by pipetting 8 uL from the normalized sub-cultured precultures to the corresponding condition of the deep well block. Base medium lacking carbohydrate was added to bring all conditions to 1200 uL. Deep well blocks were covered with a breathable seal and incubated at 37 degrees Celsius for approximately 48 hours.

Passaging of community cultures was performed by transferring 24 uL (50x) of well mixed culture to a new deep well block containing identical media. This block was incubated as previously described. The spent cultures were sampled for OD600, 16S sequencing, pH, and organic acid measurements as follows. Linear range optical density was measured by diluting 100 uL of culture in 100 uL of 1x PBS. The OD600 was calculated as 2*(Measured – Blank), where blank indicates the OD600 of 100 uL of blank media diluted in 100 uL of 1x PBS. Supernatant for organic acids and pH measurement were collected by first centrifuging 300 uL of culture at 4000 rpm for 25 minutes and removing 200 uL to a skirted PCR plate. The pH of 20 uL of supernatant was measured using a spectrophotometric phenol red assay, as described in Clark 2021. The cell pellet was washed with 500 uL of sterile 1x PBS to help remove residual fiber; centrifugation was performed at 4000 RPM for 20 minutes and supernatant was discarded. Cell pellets were stored at −80 degrees Celsius. Monoculture experiments for timeseries growth curves were performed identically to community experiments with additional sampling of timepoint OD600 and pH measurements collected at approximately 12 hour intervals.

### Genomic DNA extraction, DNA library preparation, and sequencing

DNA extraction, library preparation, and sequencing were performed according to methods described in Hromada 2021 and Clark 2021.^109,110^ Cell pellets from 300 uL of culture were washed twice with 500 uL of 1x Phoshate buffered saline to reduce residual fiber and stored at −80C following experiments. Genomic DNA was extracted using a 96-well plate adaption of the DNeasy protocol (Qiagen). Genomic DNA was normalized to 1 ng/uL in molecular grade water and stored at −20C. Dual-indexed primers for multiplexed amplicon sequencing of the v3-v4 region of the 16S gene were designed as described previously, and arrayed in 96-well, skirted PCR plates (Thomas Scientific) using an acoustic liquid handling robot (Echo LabCyte). Genomic DNA and PCR master mix were added to primer plates and amplified prior to sequencing on an Illumina MiSeq platform (Illumina).

### Organic Acids Quantification using HPLC

Supernatant samples were thawed before the addition of 2 μL of H2SO4 to precipitate any components that might be incompatible with the running buffer. The samples were then centrifuged at 2400 × g for 10 min and 150 μL of each sample was filtered through a 0.2 μm filter using a vacuum manifold and aliquoted to HPLC vials. HPLC analysis was performed using a Shimadzu HPLC system equipped with a SPD-20AV UV detector (210 nm). Compounds were separated on a 250 × 4.6 mm Rezex© ROA-Organic acid LC column (Phenomenex Torrance, CA) run with a flow rate of 0.3 mL min−1 and at a column temperature of 50 °C. The samples were held at 4 °C prior to injection. Separation was isocratic with a mobile phase of HPLC grade water acidified with 0.015 N H2SO4 (415 µL L−1). Standards were run before and after each set of 96 samples. Standards were 100, 20, and 4 mM concentrations of butyrate, lactate, and acetate. The injection volume for both samples and standards were 20 µL. The data was analyzed Shimadzu LabSolutions software package.

### Colony forming unit assay

Colony forming units were immediately measured after collecting mouse fecal samples, using an agar which supported growth of all species in monoculture (**Supplementary Data 5**). The agar was a 1:1 mixture of Anaerobe basal broth powder (Oxoid) and Reinforced clostridial medium powder (Oxoid) with 15 g/L of bacteriological agar (Dot). The powdered components were prepared at 2x concentration and sterile filtered to preserve heat-sensitive components. The agar was prepared at 2x concentration and autoclaved with a stir bar. The autoclaved and filtered portions were mixed at a 1:1 ratio prior to pouring plates. Approximately 20 mg of fecal sample was transferred to an Eppendorf tube, brought in an anaerobic chamber (Coy labs) immediately after sampling, and 1000 uL of 1x PBS was added. The fecal pellets were allowed to soak for one hour to improve resuspension. Pre-reduced agar plates were placed in the incubator at this time, to improve absorption of liquid. Pellets were resuspended by pipetting a volume of 800 uL roughly 40 times, and three 20 uL technical replicates were immediately transferred to a round bottom 96-well plate for serial dilution. Serial dilutions were performed at a 20:180 uL dilution after and mixed by pipetting 100 uL 20 times. Each dilution series was mixed by pipetting a 100 uL volume 9 times immediately prior to spotting five uL. Plates were incubated upside-down in bags to prevent drying, for 48 hours. CFU/mg was calculated as (colony count) * (overall dilution factor) / (spot volume). Three technical replicates were averaged for each of the five biological replicate mice.

### DNA extraction from fecal and cecal samples

Genomic DNA extraction from fecal and cecal samples was performed as described previously (Feng, 2022) based on (Goodman, 2011) with some modifications. Fecal samples (∼20 mg) were transferred into solvent-resistant screw-cap tubes (Sarstedt Inc) with 1.2 g of 0.1 mm zirconia/silica beads (BioSpec Products) and one 3.2 mm stainless steel bead (BioSpec Products). The samples were resuspended in 500 μL of Buffer A (200 mM NaCl (DOT Scientific), 20 mM EDTA (Sigma) and 200 mM Tris·HCl pH 8.0 (Research Products International), 210 μL 20% SDS (Alfa Aesar) and 500 μL phenol/chloroform/isoamyl alcohol (Invitrogen). Cells were lysed by mechanical disruption with a bead-beater (BioSpec Products) for 3 min twice to prevent overheating. Next, cells were centrifuged for 5 min at 8,000 x g at 4°C, and the supernatant was transferred to a Eppendorf tube. We added 60 μL 3M sodium acetate (Sigma) and 600 μL isopropanol (LabChem) to the supernatant and incubated on ice for 1 hr. Next, samples were centrifuged for 20 min at 18,000 x g at 4°C. The harvested DNA pellets were washed once with 500 μL of 100% ethanol (Koptec). The remaining trace ethanol was removed by air drying the samples. Finally, the DNA pellets were then resuspended into 200 μL of AE buffer (Qiagen). The crude DNA <u>extracts</u> were purified by a Zymo DNA Clean & Concentrator™-5 kit (Zymo Research). Library preparation and sequencing proceeded according to previous description for synthetic community experiments.

### Quantification of mouse fecal SCFA via headspace gas chromatography

Samples were resuspended by adding approximately 20 mg of fecal pellet to a water of volume (300 uL – mass of fecal pellet). Samples were soaked for one hour to facilitate resuspension and resuspended by pipetting. All materials were thoroughly chilled on ice at this time. Suspended samples were added to 20 mL headspace crimp seal vials (Restek) containing two grams of sodium bisulfate, 1 mL of freshly prepared 60 micromolar 2-butanol was added, and vials were crimp-sealed immediately. Samples were run on Shimadzu GC instrument with a Polar wax column. Data was analyzed in Shimadzu LabSolutions software.

### Bioinformatic analysis of strain abundances

Sequencing data were analyzed as described in Hromada 2021. Basespace Sequencing Hub’s FastQ Generation demultiplexed the indices and generated FastQ files. Paired reads were merged using PEAR (Paired-End reAd mergeR) v0.9.^112^ Reads were mapped to a reference database of species used in this study, using the mothur v1.40.5, and the Wang method.^113,114^ Relative abundance was calculated by dividing the read counts mapped to each organism by the total reads in the sample. Estimated absolute abundance was calculated by multiplying the relative abundance of an organism by the OD600 of the sample. Samples were excluded from further analysis if > 1% of the reads were assigned to a species not expected to be in the community (indicating contamination).

### Multidimensional scaling of the species-fiber design space

Communities can be represented as binary vectors that capture the presence or absence of individual species and fibers in the community. However, considering only the presence or absence of species and fibers, the full design space results in a Boolean hypercube, which is a perfectly symmetrical object that cannot be optimally projected into lower dimensions. By augmenting each community vector to capture information about species phylogeny, different community vectors can vary in their alignment depending on phylogenetic similarity. Using the phylogenetic tree of the 15 species considered in this study (**Fig. 1a**), we augment community vectors with 13 additional elements corresponding to the 13 parent nodes in the phylogenetic tree. Community vectors that contain a species that is a descendant of a parent node in the phylogenetic tree will populate the element of the vector corresponding to that node. We can therefore represent any species-fiber combination as a 15+6+13=34-dimensional vector that captures presence or absence of the 15 species and 6 fibers, as well as the phylogenetic information of the species. Consequently, the alignment of two vectors corresponding to two different communities depends on the phylogenetic relationships between species. To create the functional landscape of the full set of experimentally characterized communities, we represented each community as a 34-dimensional vector and performed multidimensional scaling (MDS) using a Euclidean distance metric to project the data onto 2-dimensions using Scikit-Learn’s MDS function^115^.

### Analysis of variance (ANOVA) to compare butyrate production between groups

The statistical analysis to compare butyrate production in sets of experimental conditions containing combinations of *A. caccae, B. uniformis*, and inulin was performed using one-way analysis of variance (ANOVA) and Tukey’s honest significant difference (HSD) procedure to control for family-wise type I error for all 28 pairwise comparisons between the eight groups. The statistical analysis to compare butyrate production in groups containing combinations of *A. caccae, B. uniformis*, *P. copri*, and inulin was performed using one-way analysis of variance (ANOVA) and Tukey’s honest significant difference (HSD) procedure to control for family-wise type I error for all 120 pairwise comparisons between the 16 groups.

## Data and code availability

All code and data used for model fitting and experimental design will be publicly available upon acceptance of the manuscript for publication.

## Author contributions

B.M.C., J.T., and O.S.V. conceived the study. B.M.C. carried out experiments. J.T. performed computational modeling and experimental design. J.T. and B.M.C. performed statistical analysis of experimental data. B.M.C., J.T., and O.S.V. analyzed data and wrote the paper. O.S.V. secured funding.

## Acknowledgements

The authors would like to thank Federico Rey and Eugenio I. Vivas for assistance in performing germ-free mouse experiments. This research was supported by National Institutes of Health under Grant Number R35GM124774, R01EB030340 and Army Research Office W911NF-19-1-0269.The funders had no role in the study conceptualization, data analysis, decision to publish, or preparation of the manuscript.

