## Supplementary Information for "Machine learning-guided design of human gut microbiome dynamics in response to dietary fibers"

| Model | Training data | Validation Step 1: cross validation on corresponding DTL data | Validation step 2: retroactive validation on additional DTL data |
|---|---|---|---|
| MR0 | DTL0 | Training – training sets for 20-fold cross validation on DTL0; monoculture 48hr timepoint<br><br>Test – test sets for 20-fold cross validation with DTL0 data | Training – All DTL0 data; monoculture 48hr timepoint<br><br>Test – All DTL1, 2, 3, & VAL data |
| MR1 | DTL0 & 1 | Training – training sets for 20-fold cross validation on DTL0 & 1; monoculture 48hr timepoint<br>Test – test sets for 20-fold cross validation with DTL0 & 1 data | Training – All DTL0 & 1 data; monoculture 48hr timepoint<br><br>Test – All DTL1, 2, 3, & VAL data |
| MR2 | DTL0, 1, & 2 | Training – training sets for 20-fold cross validation on DTL0, 1, & 2; monoculture 48hr timepoint<br><br>Test – test sets for 20-fold cross validation with DTL0, 1, & 2 | Training – DTL0, 1, & 2; monoculture 48hr timepoint<br><br>Test – All DTL2, 3, & VAL data |
| MR3 | DTL0, 1, 2, & 3 | Training – training sets for 20-fold cross validation on DTL0, 1, 2, & 3; monoculture 48hr timepoint<br><br>Test – test sets for 20-fold cross validation with DTL0, 1, 2, & 3 | n/a |

**Supplementary Table 1** Model iterations and training/test data set descriptions corresponding to **Fig 2**. Models were trained on community abundance and pH data for all passages. Models were trained on organic acid data for passages one and three for DTL1, 2, and 3.

| Test | Unpaired two-sample t-test adjusted for multiple comparisons using Benjamini-Hochberg procedure | | | | | | |
|---|---|---|---|---|---|---|---|
| | | Day | | | | | |
| Metric | Group | 1 | 2 | 3 | 4 | 5 | 6 |
| Shannon Diversity | 1-2 | 0.0026 | 0.25 | 0.0017 | 0.83 | 0.51 | 0.58 |
| | 1-3 | 0.0016 | 1.68E-07 | 1.40E-08 | 4.74E-05 | 1.52E-06 | 3.55E-06 |
| | 2-3 | 1.77E-05 | 1.83E-06 | 1.40E-08 | 0.00032 | 2.01E-05 | 3.79E-06 |
| | | Welch's t-test adjusted for multiple comparisons using Benjamini-Hochberg procedure | | | | | |
| Metric | Group | 1 | 2 | 3 | 4 | 5 | 6 |
| CFU | 1-2 | 0.00083 | | 0.0028 | 0.03 | 0.029 | 0.044 |
| | 1-3 | | 0.03 | 0.0028 | 0.033 | 0.029 | 0.033 |
| | 2-3 | | | 0.003 | 0.03 | 0.20 | 0.0078 |

**Supplementary Table 2** p-values for Shannon diversity and CFU multiple comparison statistics

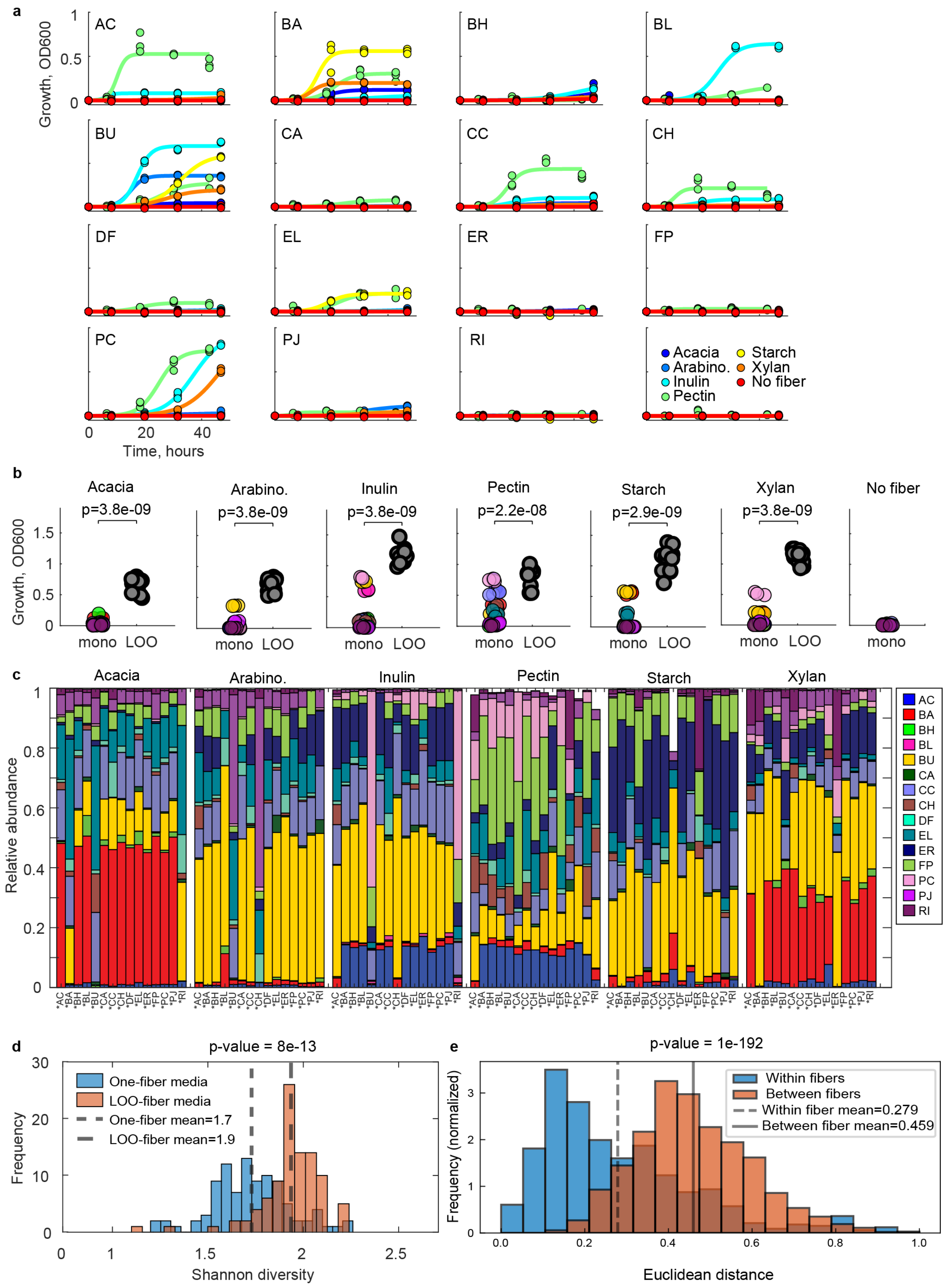
a
Growth, OD600
AC
BA
BH
BL
BU
CA
CC
CH
DF
EL
ER
FP
PC
PJ
RI
Time, hours
Acacia
Arabino.
Inulin
Pectin
Starch
Xylan
No fiber
b
Growth, OD600
Acacia
p=3.8e-09
Arabino.
p=3.8e-09
Inulin
p=3.8e-09
Pectin
p=2.2e-08
Starch
p=2.9e-09
Xylan
p=3.8e-09
No fiber
mono
LOO
c
Relative abundance
Acacia
Arabino.
Inulin
Pectin
Starch
Xylan
d
p-value = 8e-13
Frequency
One-fiber media
LOO-fiber media
One-fiber mean=1.7
LOO-fiber mean=1.9
Shannon diversity
e
p-value = 1e-192
Frequency (normalized)
Within fibers
Between fibers
Within fiber mean=0.279
Between fiber mean=0.459
Euclidean distance

**Supplementary Figure 1.** Monoculture and leave-one-out communities cultured on single-fiber media **a** Line plots indicating growth curves for all single-species, single-fiber conditions. Colored markers indicate average of n=3 biological replicates and colored lines indicate logistic model fits. Colors indicate species with the color legend shown in the bottom right panel. The bottom right panel shows timeseries sampling of blank media. **b** Categorical scatter plot of highest growth monoculture timepoint (panel a) vs. total leave-one-out (LOO) community growth in each single-fiber media. A Mann-Whitney U test is used to determine statistical significance with the p-value labeled above each plot alongside the corresponding fiber. “Acacia” indicates acacia gum and “Arabino.” indicates arabinogalactan. **c** Stacked bar plots indicating endpoint (48 hour) relative abundances of all leave-one-out communities cultured on all single-fiber media. The species not included in each community is labeled with an asterisk on the x-axis. **d** Histograms indicating distributions of Shannon diversities for single-fiber media (“one-fiber”) and leave-one-fiber-out media (five fibers) for all leave-out-out species communities. The p-value for a Mann-Whitney U test is shown above the plot. **e** Histograms of Euclidean distances between communities of leave-one-out species cultured in the same fiber (blue) and in different fibers (red-orange). The Euclidean distance between communities when cultured in the same fiber was significantly less than distances between fibers (two-tailed independent t-test p-value shown above plot)

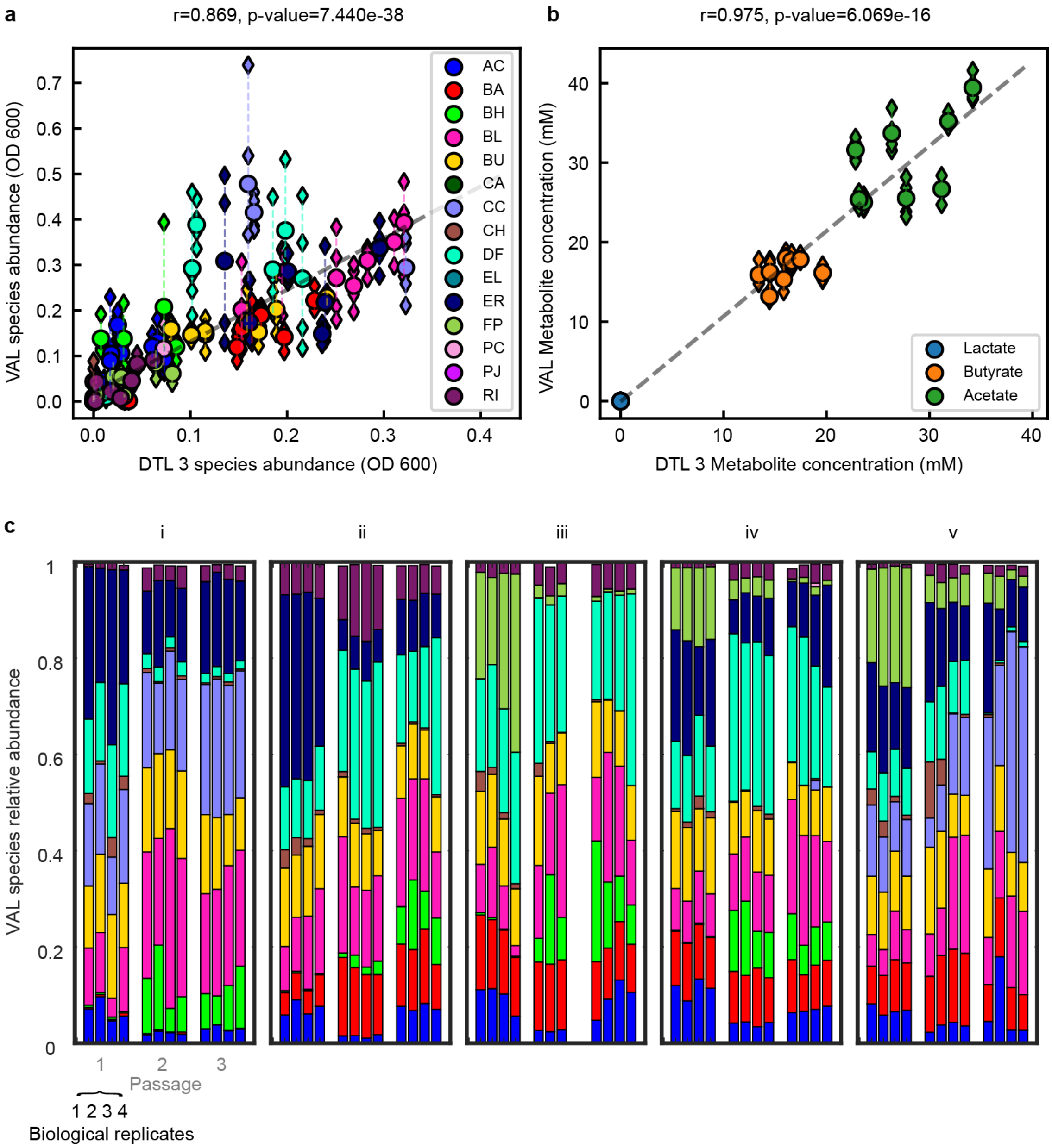

a
r=0.869, p-value=7.440e-38
VAL species abundance (OD 600)
DTL 3 species abundance (OD 600)
AC
BA
BH
BL
BU
CA
CC
CH
DF
EL
ER
FP
PC
PJ
RI
b
r=0.975, p-value=6.069e-16
VAL Metabolite concentration (mM)
DTL 3 Metabolite concentration (mM)
Lactate
Butyrate
Acetate
c
i
ii
iii
iv
v
VAL species relative abundance
1 2 3
Passage
1 2 3 4
Biological replicates

**Supplementary Figure 2.** Design objective validation data and frequency of feature selection. **a** Scatter plot of absolute abundances for the DTL3 validation experiment (n=4 biological replicates, individual replicates shown as small diamond markers and average value of replicates shown as circles) vs. abundances of (n=1 biological replicate) for corresponding communities cultured in DTL3. Dashed line indicates regression line. Pearson correlation coefficient and p-value are indicated above the plot. **b** Same as panel **a** but showing metabolite concentrations. **c** Stacked bar indicating relative abundances of all biological replicates for DTL3 validation conditions of which the mean and standard deviation are shown in Fig. 4c.

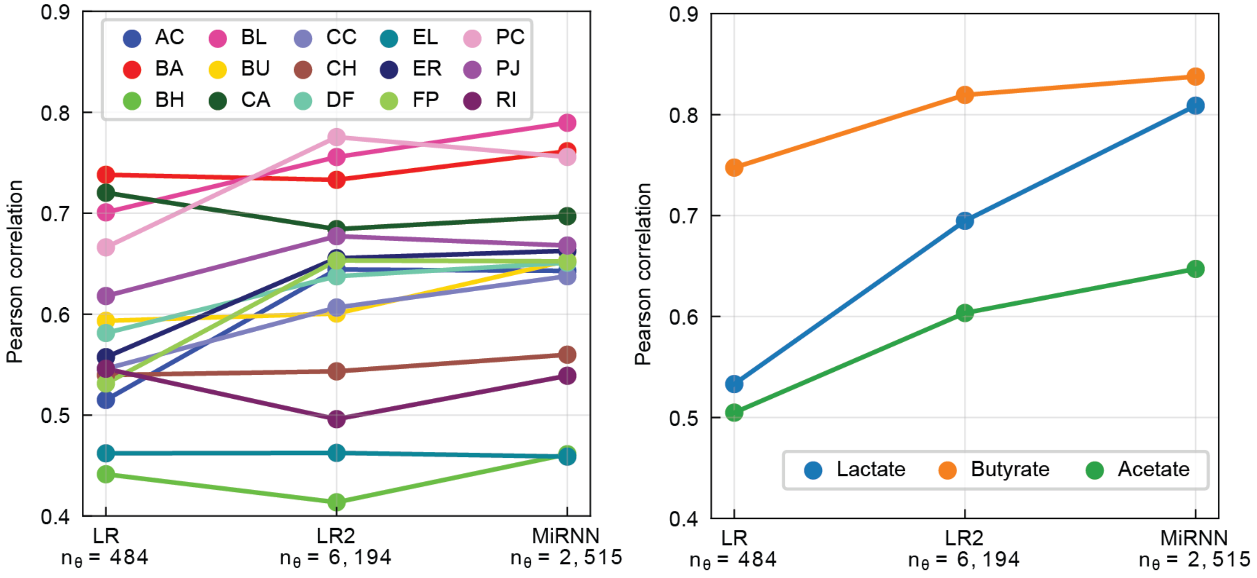

**Supplementary Figure 3.** Comparison of prediction performance between a linear state-state model with only main effects (LR), a linear state-space model with main and pairwise-effects (LR2), and the MiRNN with a 32-dimensional hidden state. The LR2 model outperformed LR1 for 11 of the 15 species and for all metabolites, and the MiRNN outperformed the LR2 model for 10 of the 15 species and all metabolites. The number of parameters of each model is shown in the x-axis, indicating that the inclusion of pairwise-terms results in more than 3 times the number of parameters of the MiRNN.

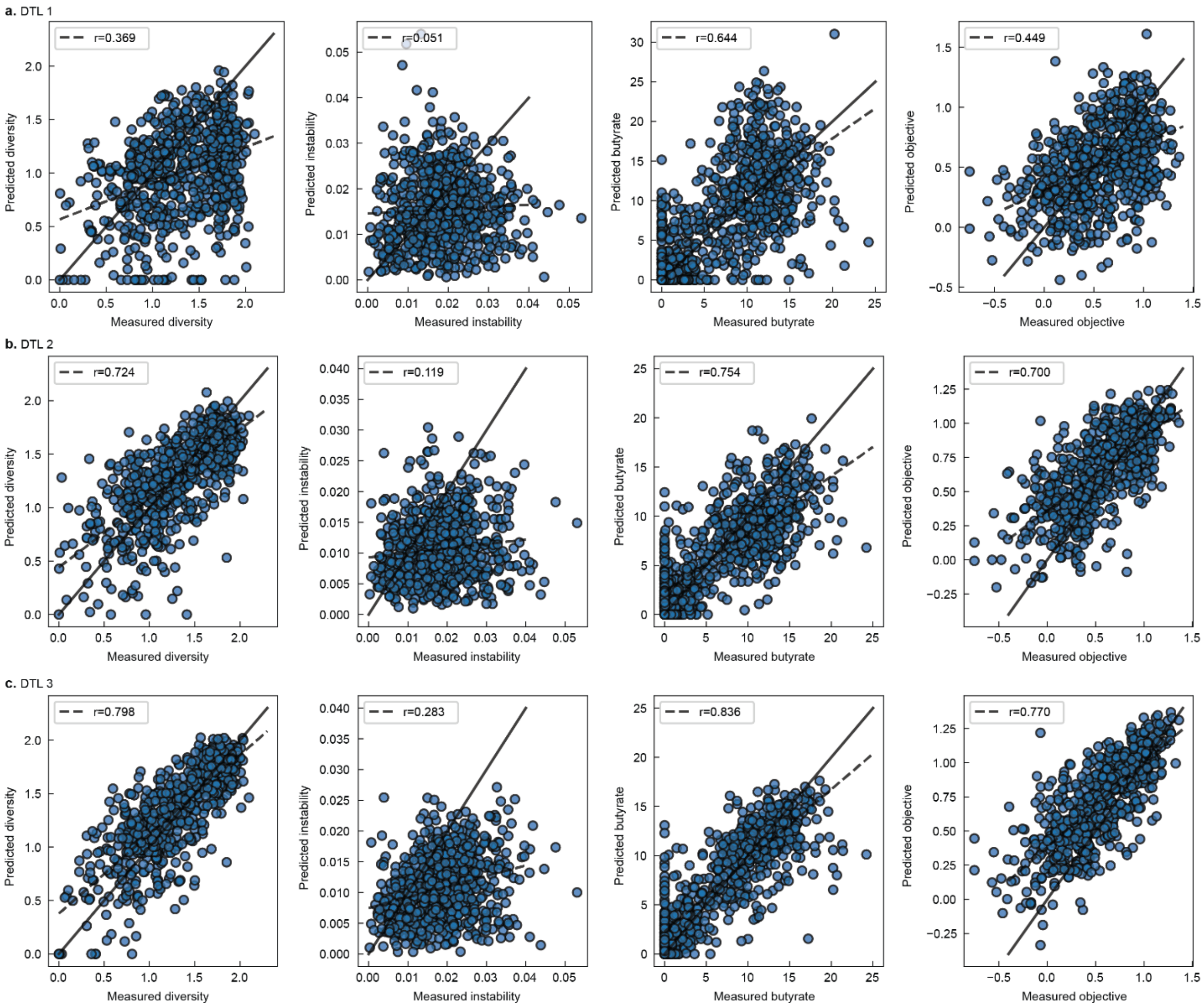

**Supplementary Figure 4.** Scatter plot of model predicted diversity, instability, butyrate, and overall objective vs. measured values. Dashed line indicates regression line. Pearson correlation corresponding to regression line are shown in top left. **a** Prediction performance on held-out data using only data from DTL 0-1 as training data. **b** Prediction performance on held-out data using only data from DTL 0-2 as training data. **c** Prediction performance on held-out data using data from DTL 0-3 as training data.

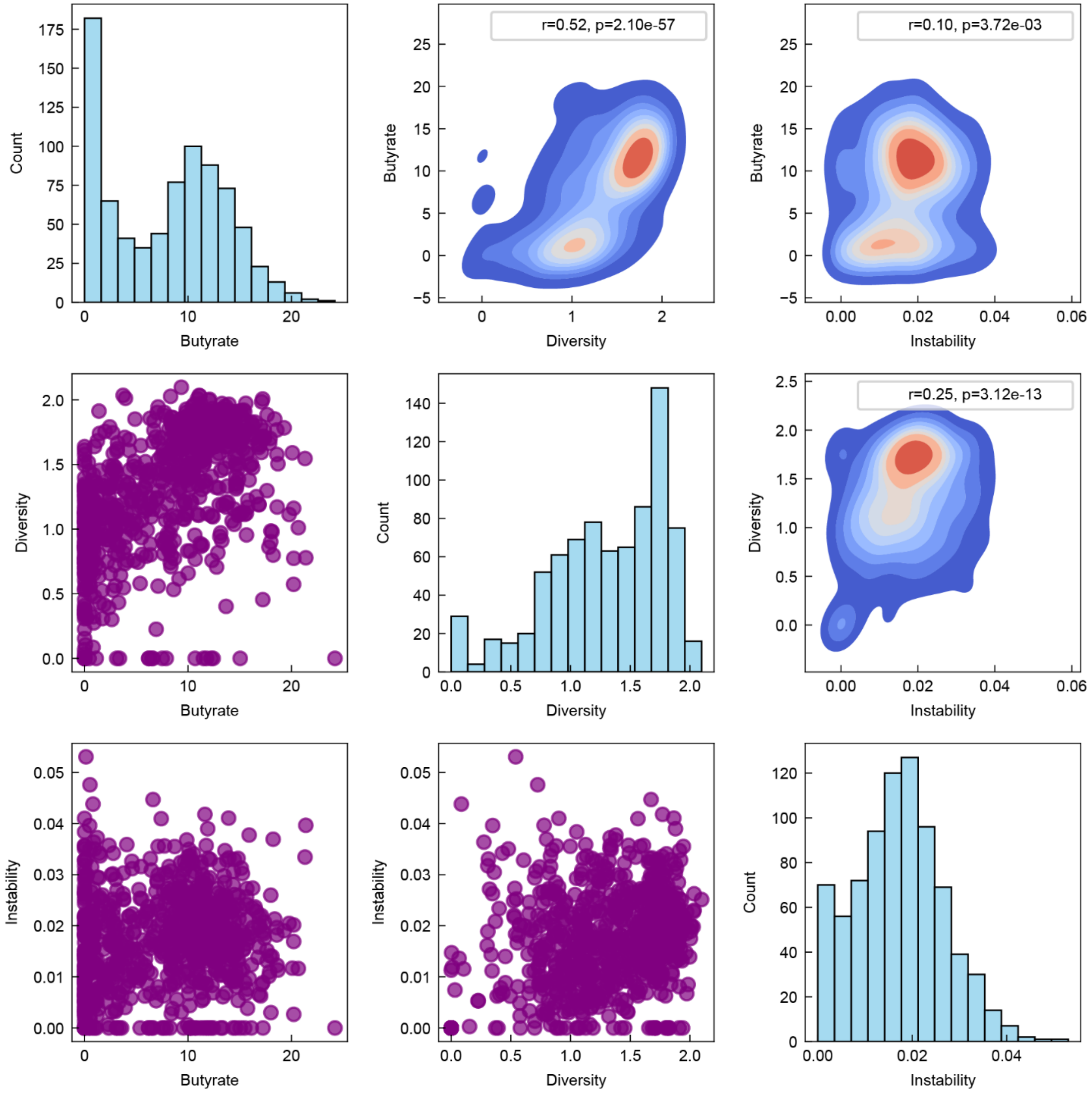

**Supplementary Figure 5.** Histograms (diagonal), scatter plots (lower triangle), and kernel density estimate plots (upper triangle) of experimentally measured butyrate, diversity, and instability.

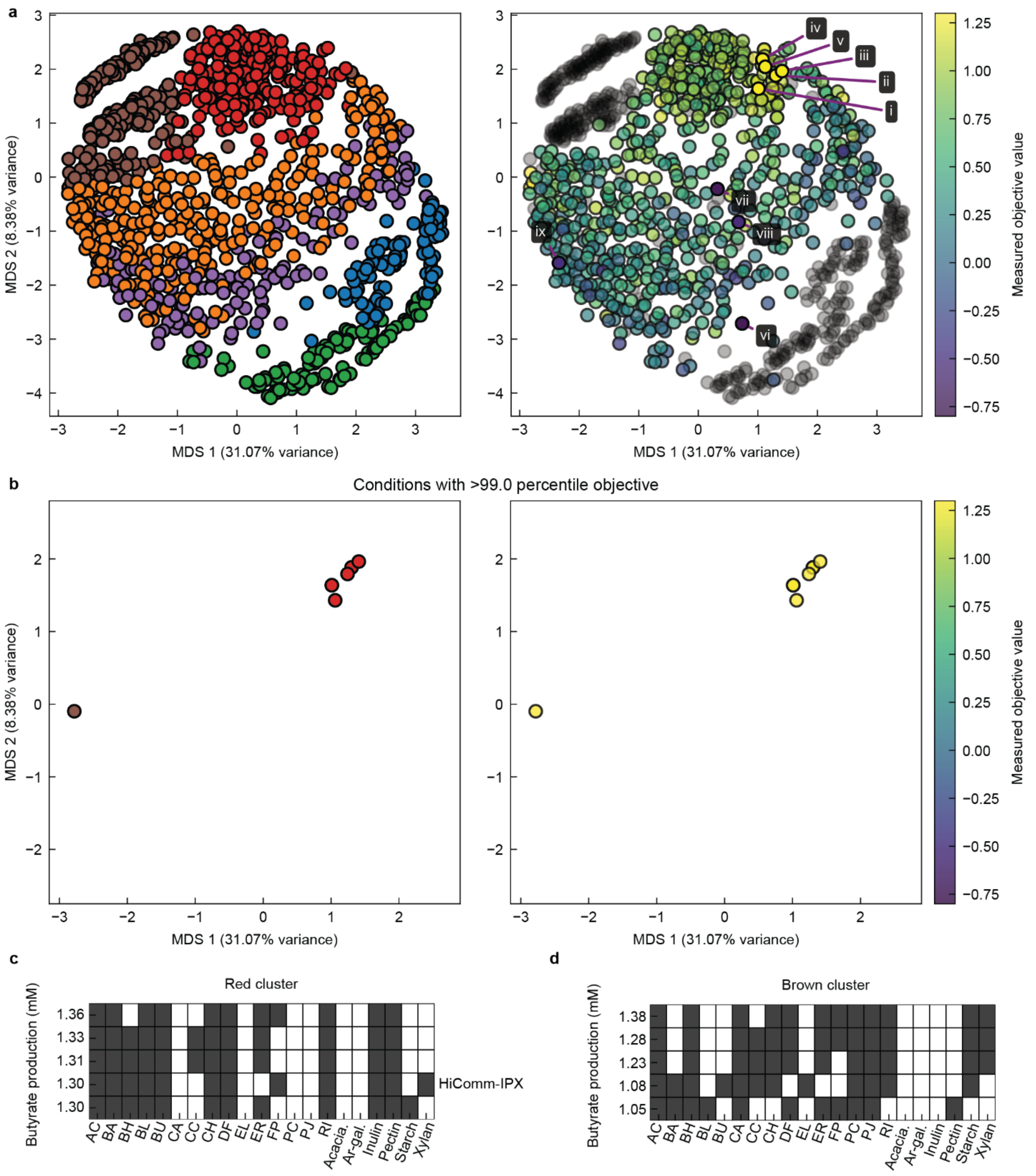

**Supplementary Figure 6. K-means clustering of the design space. a** Left panel shows K-means clustering with K=6 of the design space where each condition is colored according to its cluster assignment. Right panel shows each condition where colors represent the measured objective value. **b** Both panels show remaining conditions and cluster assignments after thresholding only conditions in the 99$^{th}$ percentile of measured objective values. **c** Presence (dark squares) or absence (blank squares) of species or fibers in the set of conditions in the red cluster with the 5 highest measured objective values. **d** Same as panel c but showing conditions from the brown cluster.

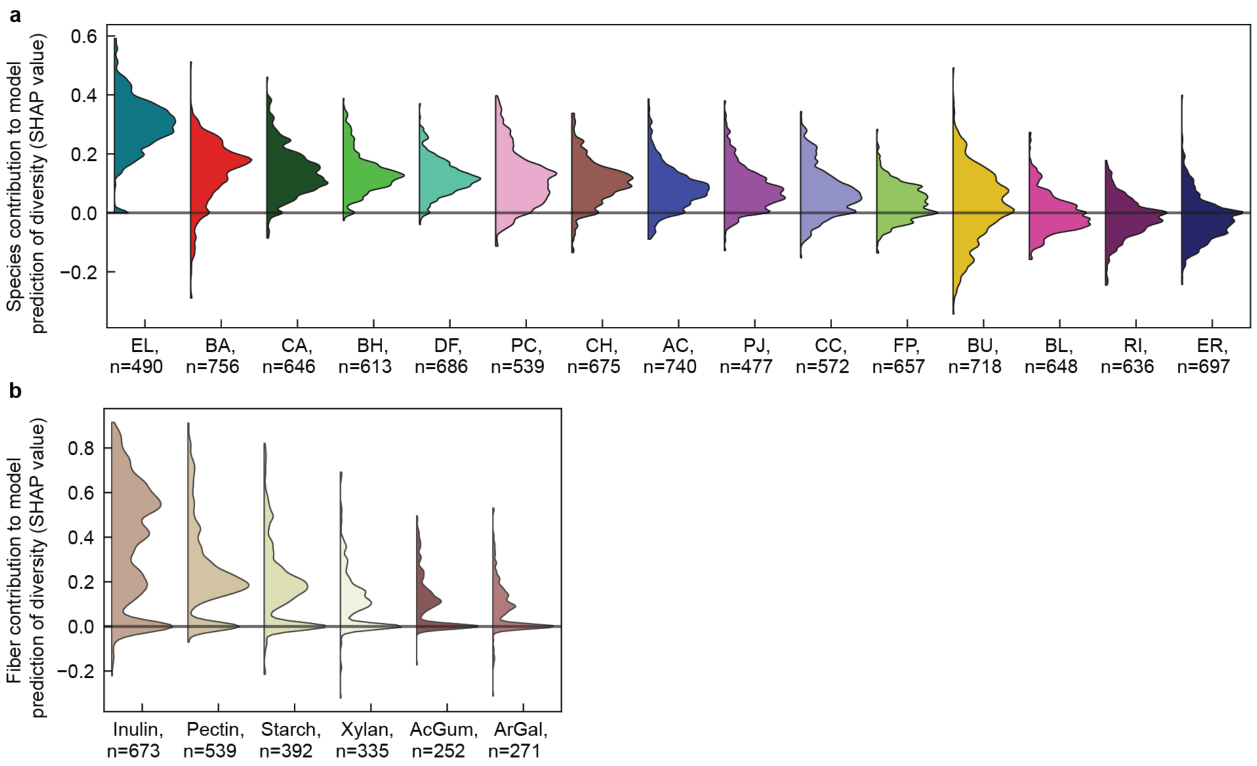

**Supplementary Figure 7. Understanding drivers of diversity using explainable machine learning. a** Distributions of SHAP values (Shapley additive explanation) of species contribution to model predicted diversity. SHAP values in the plotted distributions correspond to conditions with the species present, with the number of points in each distribution indicated in the x-axis tick labels. The horizontal line indicates a value of zero. The distributions are sorted by highest to lowest median value. **b** Distributions of SHAP values (Shapley additive explanations) of fiber contribution to model predicted diversity for conditions containing respective fiber; formatting matches panel a. Acacia gum and arabinogalactan are abbreviated as “Acacia.” and “Ar-gal.”, respectively.

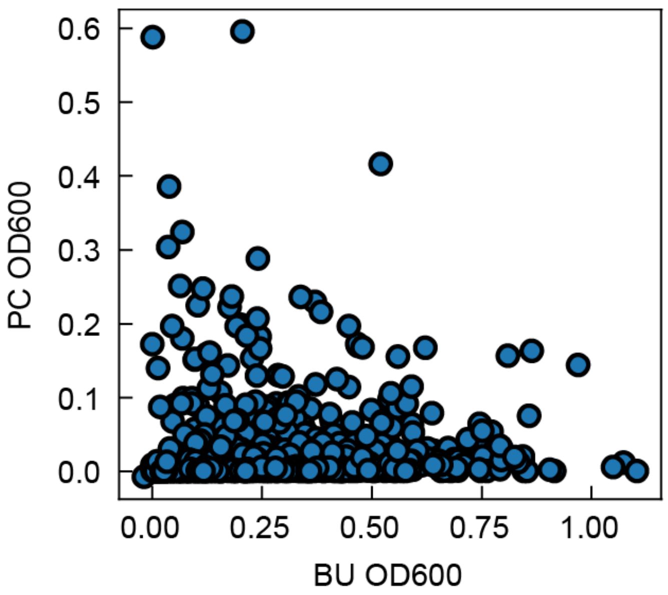

**Supplementary Figure 8. Scatter plot of the absolute abundance of *P. copri* (PC) versus *B. uniformis* (BU)** A scatter plot of experimentally measured absolute abundance (OD600) of *P. copri* versus *B. uniformis* from all experimental conditions that contained both species suggests competitive exclusion, where if one species has a high absolute abundance, the other has a low absolute abundance.

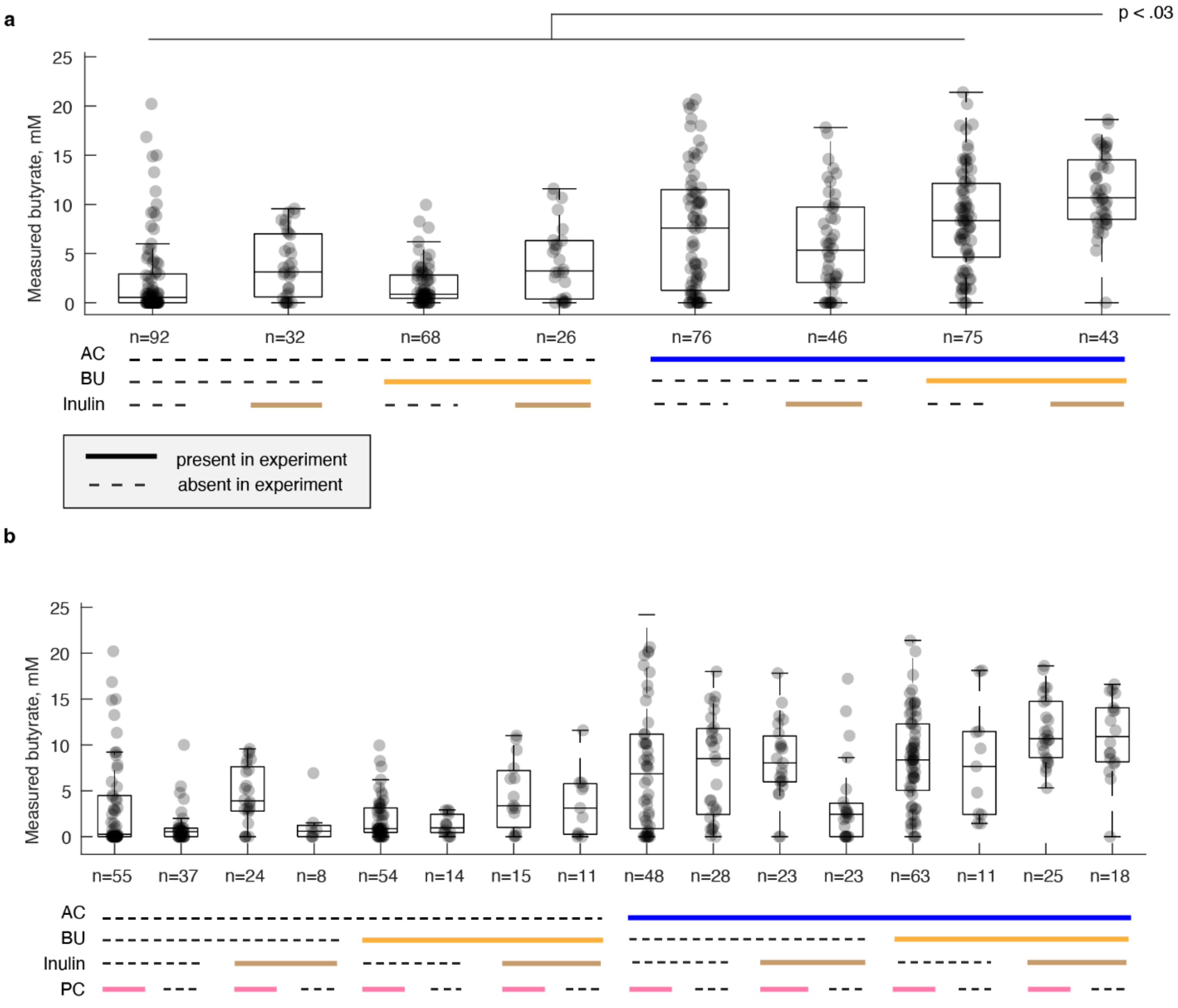

**Supplementary Figure 9. Analysis of passage three experimental butyrate measurements for higher-order interactions. a** Categorical scatter plot of experimentally measured butyrate concentrations for every combination of presence and absence of *A. caccae* (AC), *B. uniformis* (BU), and inulin for passage three. Inulin presence indicates that inulin was the only fiber present in the condition. Presence is indicated by the solid line of which the color corresponds to the variable labeled on the left, and absence is indicated by a dashed black line. Box plots indicate summary statistics of the distribution with center line, box edges, and whiskers indicating median, quartiles, and quartile plus 1.5 times interquartile range, respectively. The statistical comparison at the top of the panel indicates group eight (copresence of AC, BU, and inulin) is significantly higher than all other individual groups using pairwise comparisons (p-values < .03). All pairwise group comparisons are made using one-way analysis of variance (ANOVA) followed by Tukey's honest significant difference (HSD) procedure using MATLAB's "anova1" and "multcompare" functions. All 28 pairwise comparisons are reported in Supplemental Data. **b** Categorical scatter plot of experimentally measured butyrate concentrations for every combination of presence and absence of *A. caccae* (AC), *B. uniformis* (BU), inulin, and *P. copri* (PC) for passage one. All 120 pairwise group comparisons are made using one-way analysis of variance (ANOVA) followed by Tukey's honest significant difference (HSD) procedure using MATLAB's "anova1" and "multcompare" functions and are reported in Supplemental Data.

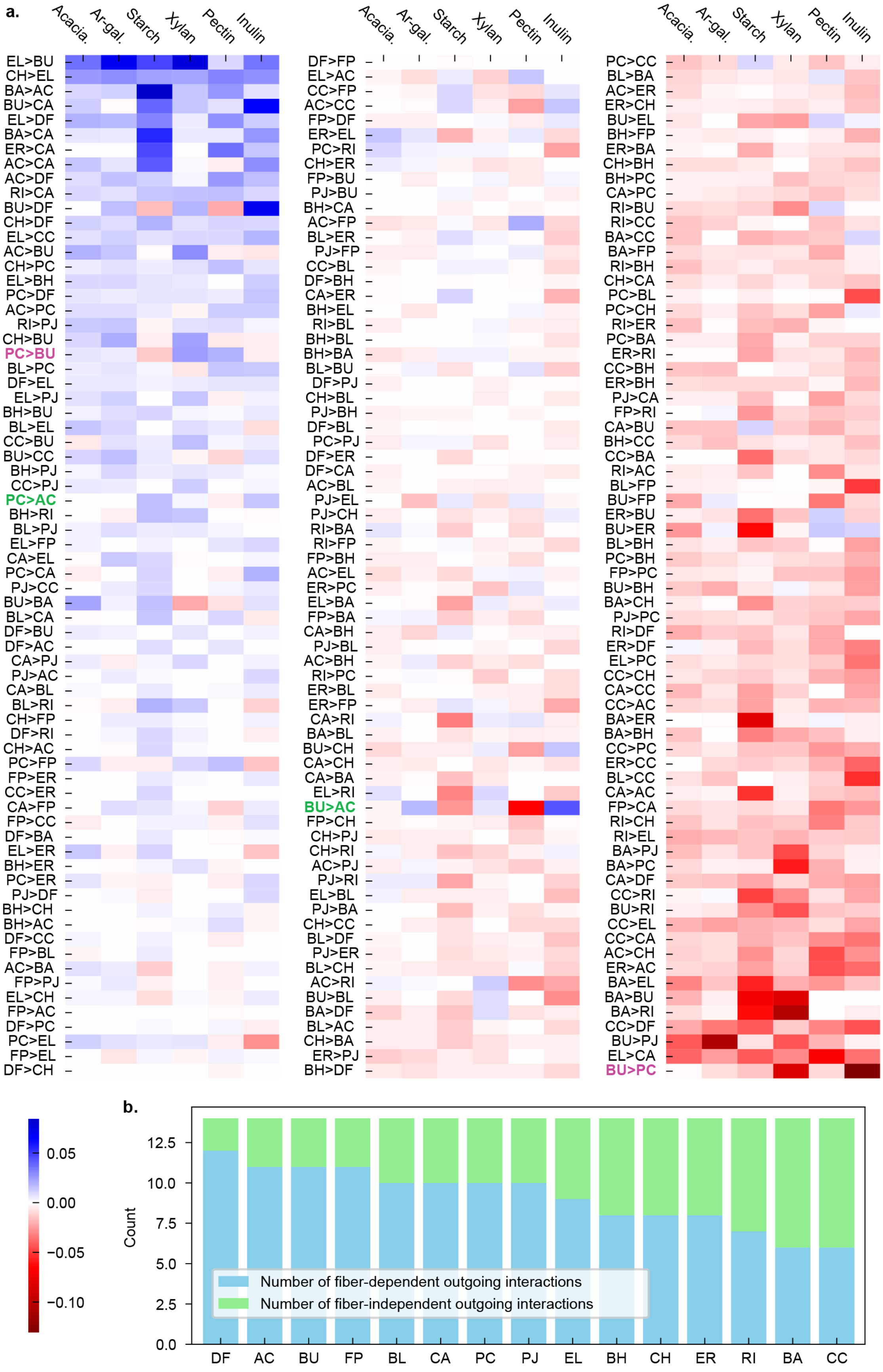
a.
Acacia.
Ar-gal.
Starch
Xylan
Pectin
Inulin
EL>BU
CH>EL
BA>AC
BU>CA
EL>DF
BA>CA
ER>CA
AC>CA
AC>DF
RI>CA
BU>DF
CH>DF
EL>CC
AC>BU
CH>PC
EL>BH
PC>DF
AC>PC
RI>PJ
CH>BU
PC>BU
BL>PC
DF>EL
EL>PJ
BH>BU
BL>EL
CC>BU
BU>CC
BH>PJ
CC>PJ
PC>AC
BH>RI
BL>PJ
EL>FP
CA>EL
PC>CA
PJ>CC
BU>BA
BL>CA
DF>BU
DF>AC
CA>PJ
PJ>AC
CA>BL
BL>RI
CH>FP
DF>RI
CH>AC
PC>FP
FP>ER
CC>ER
CA>FP
FP>CC
DF>BA
EL>ER
BH>ER
PC>ER
PJ>DF
BH>CH
BH>AC
DF>CC
FP>BL
AC>BA
FP>PJ
EL>CH
FP>AC
DF>PC
PC>EL
FP>EL
DF>CH
DF>FP
EL>AC
CC>FP
AC>CC
FP>DF
ER>EL
PC>RI
CH>ER
FP>BU
PJ>BU
BH>CA
AC>FP
BL>ER
PJ>FP
CC>BL
DF>BH
CA>ER
BH>EL
RI>BL
BH>BL
BH>BA
BL>BU
DF>PJ
CH>BL
PJ>BH
DF>BL
PC>PJ
DF>ER
DF>CA
AC>BL
PJ>EL
PJ>CH
RI>BA
RI>FP
FP>BH
AC>EL
ER>PC
EL>BA
FP>BA
CA>BH
PJ>BL
AC>BH
RI>PC
ER>BL
ER>FP
CA>RI
BA>BL
BU>CH
CA>CH
CA>BA
EL>RI
BU>AC
FP>CH
CH>PJ
CH>RI
AC>PJ
PJ>RI
EL>BL
PJ>BA
CH>CC
BL>DF
PJ>ER
BL>CH
AC>RI
BU>BL
BA>DF
BL>AC
CH>BA
ER>PJ
BH>DF
PC>CC
BL>BA
AC>ER
ER>CH
BU>EL
BH>FP
ER>BA
CH>BH
BH>PC
CA>PC
RI>BU
RI>CC
BA>CC
BA>FP
RI>BH
CH>CA
PC>BL
PC>CH
RI>ER
PC>BA
ER>RI
CC>BH
ER>BH
PJ>CA
FP>RI
CA>BU
BH>CC
CC>BA
RI>AC
BL>FP
BU>FP
ER>BU
BU>ER
BL>BH
PC>BH
FP>PC
BU>BH
BA>CH
PJ>PC
RI>DF
ER>DF
EL>PC
CC>CH
CA>CC
CC>AC
BA>ER
BA>BH
CC>PC
ER>CC
BL>CC
CA>AC
FP>CA
RI>CH
RI>EL
BA>PJ
BA>PC
CA>DF
CC>RI
BU>RI
CC>EL
CC>CA
AC>CH
ER>AC
BA>EL
BA>BU
BA>RI
CC>DF
BU>PJ
EL>CA
BU>PC
0.05
0.00
−0.05
−0.10
b.
Count
0.0
2.5
5.0
7.5
10.0
12.5
Number of fiber-dependent outgoing interactions
Number of fiber-independent outgoing interactions
DF
AC
BU
FP
BL
CA
PC
PJ
EL
BH
CH
ER
RI
BA
CC

**Supplementary Figure 10. Species interactions determined using explainable machine learning.** **a** Interspecies interactions were determined by computing the average SHAP contribution of each inoculated species to the prediction of species in the third passage. For each fiber, the average was computed over all experimentally measured conditions that included only that fiber. Interactions hypothesized to promote butyrate production are highlighted in green and interactions hypothesized to inhibit butyrate production are highlighted purple. **b** Each of the 15 species has 14 outgoing interactions with other species. Of the 14 outgoing interactions, the stacked bar plot shows how the interactions partitioned into those that remained either positive or negative in all fiber conditions (green) or changed in sign depending on the fiber (blue).

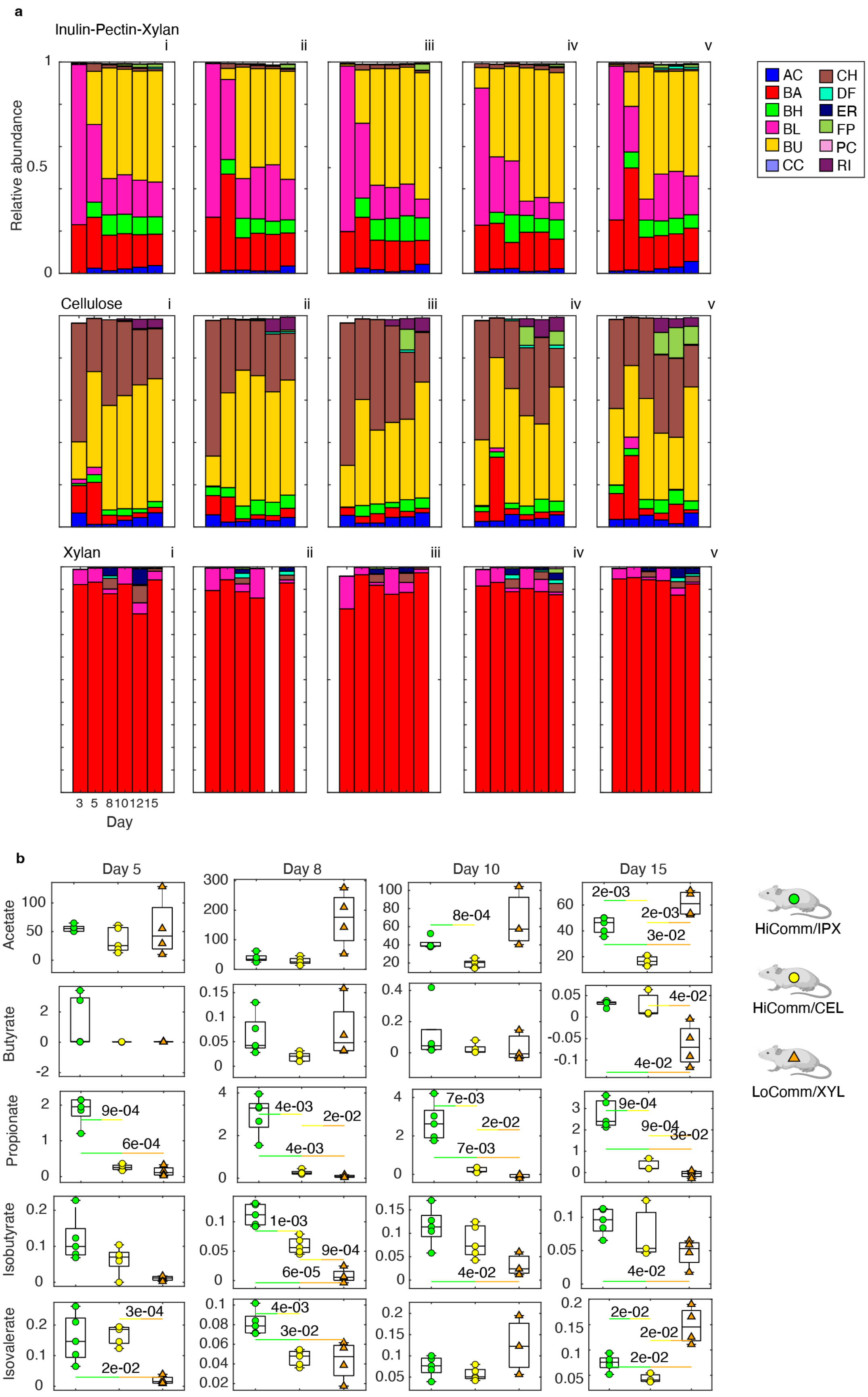

a
Inulin-Pectin-Xylan
i
ii
iii
iv
v
Relative abundance
1
0.5
0
Cellulose
Xylan
3 5 8101215
Day
AC
BA
BH
BL
BU
CC
CH
DF
ER
FP
PC
RI
b
Day 5
Day 8
Day 10
Day 15
Acetate
Butyrate
Propionate
Isobutyrate
Isovalerate
HiComm/IPX
HiComm/CEL
LoComm/XYL

**Supplementary Figure 11. Biological replicates of species abundance and short-chain fatty acids (SCFA) for gnotobiotic mouse model. a** Stacked bar plots of community composition where height indicates relative abundance of species for n=5 biological replicates labeled i-v. Colors indicate species according to legend in the lower righthand portion of **Fig. 6f**. **b** Categorical scatter plots of fecal short chain fatty acid (SCFA). Markers indicate the average concentration of SCFA for a given mouse over all available samples. Box plots indicate summary statistics for n=5 replicates where middle line indicates median, box edges indicate quartiles, and whiskers indicate maximum and minimum datapoints. All p-values < 0.05 for Welch's t-test adjusted for multiple comparisons via Benjamini-Hochberg procedure are indicated in the plot. Marker style and color is indicated in **Fig. 6a** a legend.
